# Integration of prophages into CRISPR loci remodels viral immunity in *Streptococcus pyogenes*

**DOI:** 10.1101/2020.10.09.333658

**Authors:** Andrew Varble, Edmondo Campisi, Chad W. Euler, Jessica Fyodorova, Jakob T Rostøl, Vincent A. Fischetti, Luciano A. Marraffini

## Abstract

CRISPR loci are composed of short DNA repeats separated by sequences that match the genomes of phages and plasmids, known as spacers. Spacers are transcribed and processed to generate RNA guides used by CRISPR-associated nucleases to recognize and destroy the complementary nucleic acids of invaders. To counteract this defense, phages can produce small proteins that inhibit these nucleases. Here we demonstrate that the ΦAP1.1 temperate phage utilizes an alternate approach to antagonize the type II-A CRISPR response in *Streptococcus pyogenes*. Immediately after infection this phage expresses a canonical anti-CRISPR, AcrIIA23 that prevents Cas9 function, allowing ΦAP1.1 to integrate into the direct repeats of the CRISPR locus and neutralizing immunity. However, *acrIIA23* is not transcribed during lysogeny and phage integration/excision cycles can result in the deletion and/or transfer of spacers, enabling a complex modulation of the type II-A CRISPR immune response.

## Introduction

Clustered, regularly interspaced, short, palindromic repeat (CRISPR) loci provide prokaryotic organisms with adaptive immunity against invading nucleic acids, including viruses (Barrangou et al., 2007) and plasmids (Marraffini and Sontheimer, 2008). CRISPR loci are composed of short direct repeats (DRs) separated by equally short sequences that match the genomes of the invader, known as spacers. Usually, the terminal repeat (DRt) is not identical to the rest of the DRs, and marks the end of the CRISPR array (Jansen et al., 2002). In the first phase of CRISPR immunity, new spacer sequences are acquired from the infecting DNA, and integrated between two repeats in the CRISPR array (Barrangou et al., 2007). In the second phase, spacers are transcribed and processed to generate RNA guides used by CRISPR-associated (Cas) nucleases to recognize and destroy the complementary nucleic acids of invaders (Brouns et al., 2008; Garneau et al., 2010; Marraffini and Sontheimer, 2008). Depending on the content of CRISPR-associated (*cas*) genes, CRISPR loci are classified into six different types and multiple subtypes (Makarova et al., 2020). The type II-A CRISPR-Cas system of *Streptococcus pyogenes* has been extensively studied, not only for its role in phage defense (Heler et al., 2015; Modell et al., 2017), but also for the biotechnological applications of its signature RNA-guided nuclease, Cas9 (Cong et al., 2013; Jinek et al., 2012; Mali et al., 2013). During phage infection a small number of cells acquire a 30-nucleotide spacer sequence from the viral genome, which is integrated between the 36-nucleotide DRs (McGinn and Marraffini, 2016; Wright and Doudna, 2016; Xiao et al., 2017). DRs and spacers are transcribed into a single pre-CRISPR-RNA (pre-crRNA), which is subsequently processed into mature crRNA guides after interaction of the DR and 5’ end of the trans-activating crRNA (tracrRNA), a small RNA cofactor of Cas9 (Deltcheva et al., 2011). The dsRNA formed through this interaction is cleaved by RNase III (Deltcheva et al., 2011), to create the mature crRNA and activate the Cas9 nuclease complex, which can use the spacer sequence to find complimentary targets in invading DNA (Jinek et al., 2012).

*S. pyogenes* (also known as the group A streptococcus), an important human pathogen, is the causative agent of a variety of diseases such as scarlet fever, pharyngitis, impetigo, cellulitis, necrotizing fasciitis and toxic shock syndrome (2016). A significant number of the virulence genes responsible for *S. pyogenes* pathogenicity are harbored within prophages integrated throughout the chromosome (Fischetti, 2007). Integration typically occurs after the injection and circularization of the of the viral genome, during the lysogenic cycle of temperate phages (Howard-Varona et al., 2017). It usually requires recombination between specific sequences in the phage (*attP*) and the bacterial (*attB*) DNA, which generates two new sites at the left (*attL*) and right (*attR*) borders of the inserted prophage. Changes in the environment causing bacterial stress can induce the lytic cycle of the prophage, resulting in the excision and replication of the viral genome, followed by the formation and release of new viral particles that can infect other hosts (Howard-Varona et al., 2017). In *S. pyogenes* this process contributes to the spread of the virulence genes carried by the temperate phages (Fischetti, 2007). In addition, inaccurate prophage excision and/or DNA packaging during induction can lead to the transduction of bacterial sequences into new hosts (McCullor et al., 2018). Recently, a comprehensive study of 118 complete *S. pyogenes* genome sequences determined that they collectively harbor 373 prophages (Yamada et al., 2019). In addition to the type II-A CRISPR system described above, which is present in 21 genomes, 20 *S. pyogenes* strains contained type I-C CRISPR loci, and 39 strains displayed both systems. However, genomes hosting a lower number of prophages had on average a significantly higher number of type II-A rather than type I-C spacers, suggesting that type I-C CRISPR systems are less efficient at preventing prophage spread, perhaps even non-functional (Yamada et al., 2019). The analyzed type II-A CRISPR arrays contained a total of 139 unique spacer sequences, of which 126 matched diverse *S. pyogenes* prophages and phages. Given the importance of prophage acquisition for *S. pyogenes* pathogenesis, the targeting of these mobile elements by type II-A CRISPR-Cas systems is expected to have profound consequences for the virulence of this organism.

Invaders must develop strategies to combat their host’s defenses. Type II-A CRISPR-Cas immunity is most commonly evaded by phages containing target mutations that prevent recognition and cleavage by Cas9 (Deveau et al., 2008; Gasiunas et al., 2012; Jinek et al., 2012). In addition, some phages also produce anti-CRISPR (Acr) proteins that specifically inhibit Cas nucleases (Davidson et al., 2020). Although none have been isolated from *S. pyogenes* phages, multiple Acrs that prevent the function of different type II-A Cas9 nucleases with diverse mechanisms of inhibition have been reported (Eitzinger et al., 2020; Forsberg et al., 2019; Hynes et al., 2017; Mahendra et al., 2020; Rauch et al., 2017; Uribe et al., 2019). Acrs are typically present in temperate phages and are robustly transcribed in both the lytic and latent cycles immediately after infection (Bondy-Denomy et al., 2013; Stanley et al., 2019). However, this elevated expression is not sufficient to inhibit all the crRNA-guided Cas nucleases present in the host cell and multiple rounds of infection (and Acr delivery) are required to neutralize the CRISPR immune response (Borges et al., 2018; Landsberger et al., 2018). Importantly, the initial high levels of transcription of the *acr* operon are detrimental to the propagation of the phage, as it interferes with the proper transcription of other viral genes required for the later stages of the infection cycle. This is prevented by the *aca* repressor, which is usually located next to the *acr* genes, and contains a helix-turn-helix DNA-binding domain to regulate the transcription of the operon (Stanley et al., 2019). Aca repression, however, is not complete since prophages can produce basal amounts of Acr that can inhibit CRISPR immunity in the lysogen.

In addition to the expression of virulence factors and anti-CRISPRs, temperate phages can modulate host phenotypes by integrating and disrupting genes that carry *attB* sites (Coleman et al., 1991; Lee and Iandolo, 1986; Taylor, 1963). A bioinformatic study reported two closely related *S. pyogenes* strains (SSI-1 and MGAS315) harboring prophages inserted into the repeats of the CRISPR array (Nozawa et al., 2011), an observation that suggests a new mechanism used by phages to inhibit CRISPR. Here we studied a member of this temperate phage family, ΦAP1.1, to determine whether and how type II-A CRISPR-Cas immunity is affected by prophage integration into the CRISPR repeats. We found that immediately after infection this phage expresses a canonical anti-CRISPR, AcrIIA23 that inhibits Cas9. However, *acrIIA23* is not transcribed from the prophage genome during lysogeny. Instead, the genesis of mature crRNA guides, and thus the ability of Cas9 to recognize the phage, is prevented by the integration of ΦAP1.1 into the direct repeats of the CRISPR locus. In addition, phage integration/excision cycles can result not only in the deletion of spacers, but also in the transduction of spacer-repeat units carried within the viral genomes. Our results reveal a new phage strategy that both modulates the type II-A CRISPR immune response and drives evolution on both sides of the host/pathogen conflict.

## Results

### The temperate phage ΦAP1.1 integrates into the repeat sequences of the *S. pyogenes* type II-A CRISPR array

We performed a bioinformatic search within all available *S. pyogenes* genomes to identify prophages integrated into the different CRISPR arrays carried by this organism. In agreement with previous analysis (Nozawa et al., 2011; Yamada et al., 2019), we found that while none of the type I-C loci contained prophages, twelve different strains harbored almost identical prophages inserted into the type II-A CRISPR array (Fig. S1A). In most cases only a single degenerate terminal repeat (DRt) was present and was interrupted by the viral genome, with the exception of strains 1085 and MGAS10786, where the prophage integrated into the first repeat and was followed by an intact CRISPR array.

To understand if and how temperate phages integrate into the type II-A CRISPR locus, we isolated the phage ΦAP1.1 from strain AP1 (Fiebig et al., 2015) (Fig. S1B). Sequencing of the isolated phage revealed that it is a *cos* phage harboring an *attP* site with 8 base-pairs identical to the DR and DRt, the *attB* for this phage (Fig. 1A). To facilitate the isolation and quantification of lysogens, we added a spectinomycin resistance cassette to ΦAP1.1 (Fig. S1B). This phage was then used to infect *S. pyogenes* K56 (Kjems, 1958), a strain containing a CRISPR array of 2 spacers, 2 DRs, and one DRt (Fig. 1B). In principle, integration could occur in each of the three *attB* sites contained within the repeats (DR1, DR2 and DRt), therefore we extracted genomic DNA from bacteria collected 24 hours after infection and performed PCR with primers that specifically amplify the *attL* and *attR*. We obtained three PCR products separated by approximately 66 base-pairs, indicative of phage integration into all the repeats, with a preference for DR1, DR2 and then DRt, according to the intensity of the DNA band (Fig. 1C). To confirm and quantify the integration events, we isolated 100 individual lysogens and sequenced the *attL* and *attR*. We found that indeed, ΦAP1.1 integrated into every direct repeat, but with a marked preference for DR1, where a majority of the *attL* and *attR* (Fig. 1D) sites were positioned. Due to this, prophage integration typically conserved the original repeat-spacer content, with 75/100 lysogens maintaining the two original spacers (Fig. 1E) where the *attL* and *attR* are inserted into the same repeat (Fig. 1F). Finally, a fraction of the insertion events led to CRISPR rearrangements that resulted in either the deletion of one or both spacers (23/100, Fig. 1E) where the *attL* and *attR* were located across different repeats (Fig. 1F), or in spacer duplication events, (2/100, Figs. 1E, S1C). These data demonstrate that although ΦAP1.1-like phages are typically detected within a single DRt, they are able to integrate into any repeat of a canonical CRISPR array.

**Figure 1.**
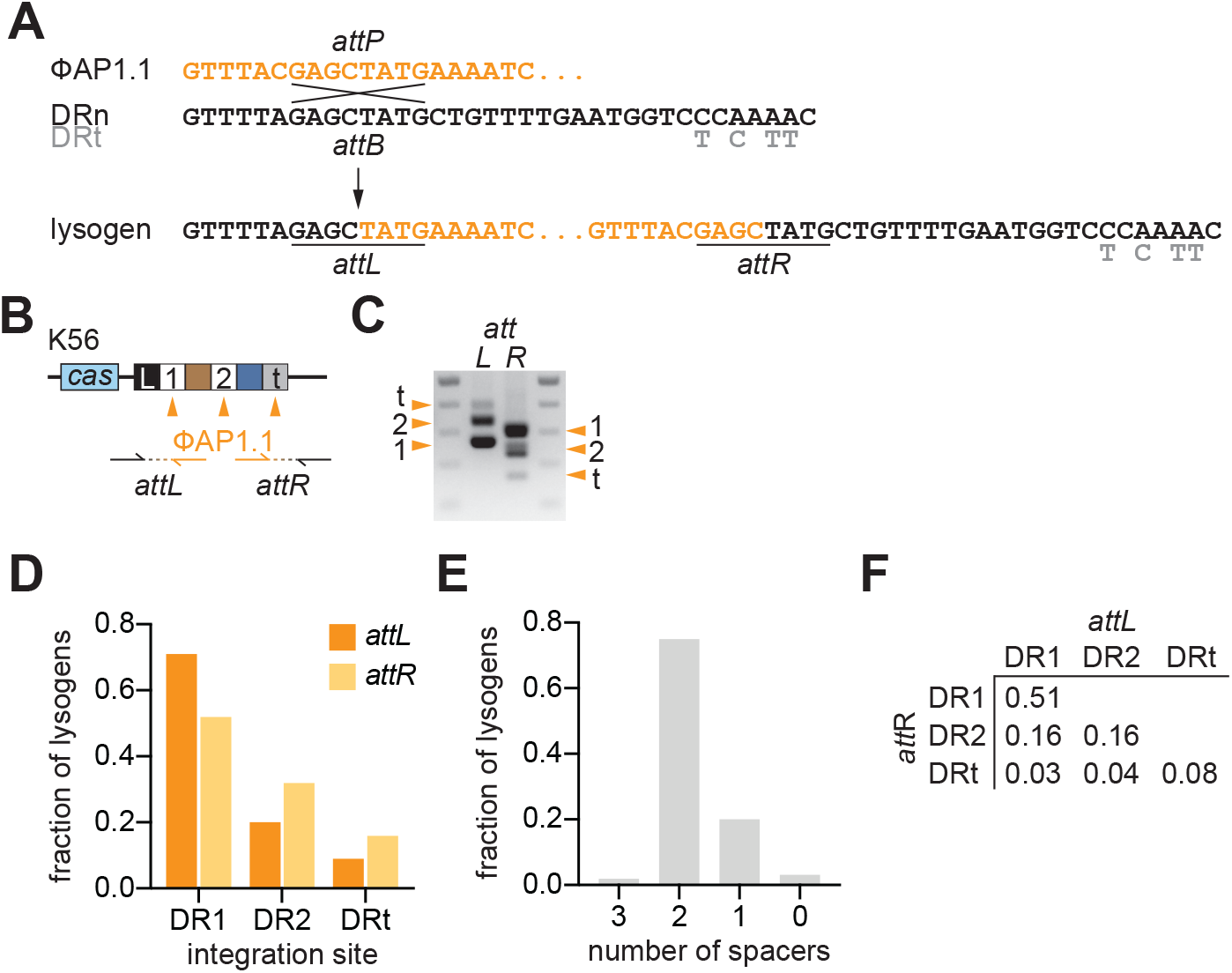
ΦAP1.1 lysogenization into the CRISPR locus. (**A**) Recombination between the *attP* site of ΦAP1.1 (orange) and the *attB* site present in the direct repeat (DR, black) of *S. pyogenes* type II-A CRISPR loci. The mutations of the terminal repeat (DRt) are shown in grey. Recombination results in prophage integration and the generation of *attL* and *attR* sites that will recombine during prophage induction and excision. (**B**) Type II-A CRISPR-*cas* locus of *S. pyogenes* K56, showing DR1 and DR2 and DRt as white or grey boxes; spacers as colored boxes. Orange arrowheads indicate the three possible integration sites of ΦAP1.1 in the different repeats. Arrows indicate the primers that anneal on the host (black) or phage (orange) genome to amplify the resulting the *attL* or *attR* sites. (**C**) Agarose gel electrophoresis of the *attL* or *attR* PCR products obtained after the amplification shown in (**B**). Orange arrowheads indicate the inferred integration sites. (**D**) Frequency of lysogens (n=100) carrying *attL* or *attR* sites in each of the DRs of the K56 CRISPR locus. (**E**) Frequency of lysogens (n=100) carrying different number of spacers. (**F**) Same as (**D**) but showing the combined distribution of *attL* or *attR* sites.

### ΦAP1.1 integration disrupts spacer transcription and crRNA processing

To determine if lysogenization could be possible in the presence of a spacer that targets the infecting phage, we investigated lysogenization in *S. pyogenes* CEM1ΔΦ (Euler et al., 2016), an SF370 (Desiere et al., 2001) derivative lacking all prophages (Fig. 2A). In CEM1ΔΦ, one of its six spacers (*spc4*) has a partial match to ΦAP1.1 (Fig. S1B, D). We transformed this strain and a mutant lacking spacers 2-5 (CEM1ΔΦ[Δ2-5]) with a plasmid harboring the phage target (pTgt4) and confirmed that this sequence is targeted by the type II-A CRISPR-Cas system (Fig. 2B). We then infected strain CEM1ΔΦ with ΦAP1.1 and analyzed the location of *attL* and *attR* by sequencing the PCR products of 100 lysogens. Interestingly, integration of the *attL* was only observed in the first five DRs, but not in DR6 nor DRt, of the CRISPR array of strain CEM1ΔΦ (Fig. 2C). Other parameters were similar to those obtained for strain K56: *attR*s were detected in every DR (Fig. 2C) and most integration events maintained the original composition of the CRISPR array, with a low frequency of spacer deletions (Figs. 2D, E). The presence of *spc4* in most lysogens suggested that the immunity mediated by this spacer was neutralized by the insertion of ΦAP1.1 into the CRISPR array. Indeed, targeting of the pTgt4 plasmid was abolished in lysogens that had the prophage integrated into DRs one through five (maintaining the entire spacer repertoire), as well as a lysogen where all spacers were eliminated due to integration into both DR1 and DRt (DR1-t, Fig. 2F).

**Figure 2.**
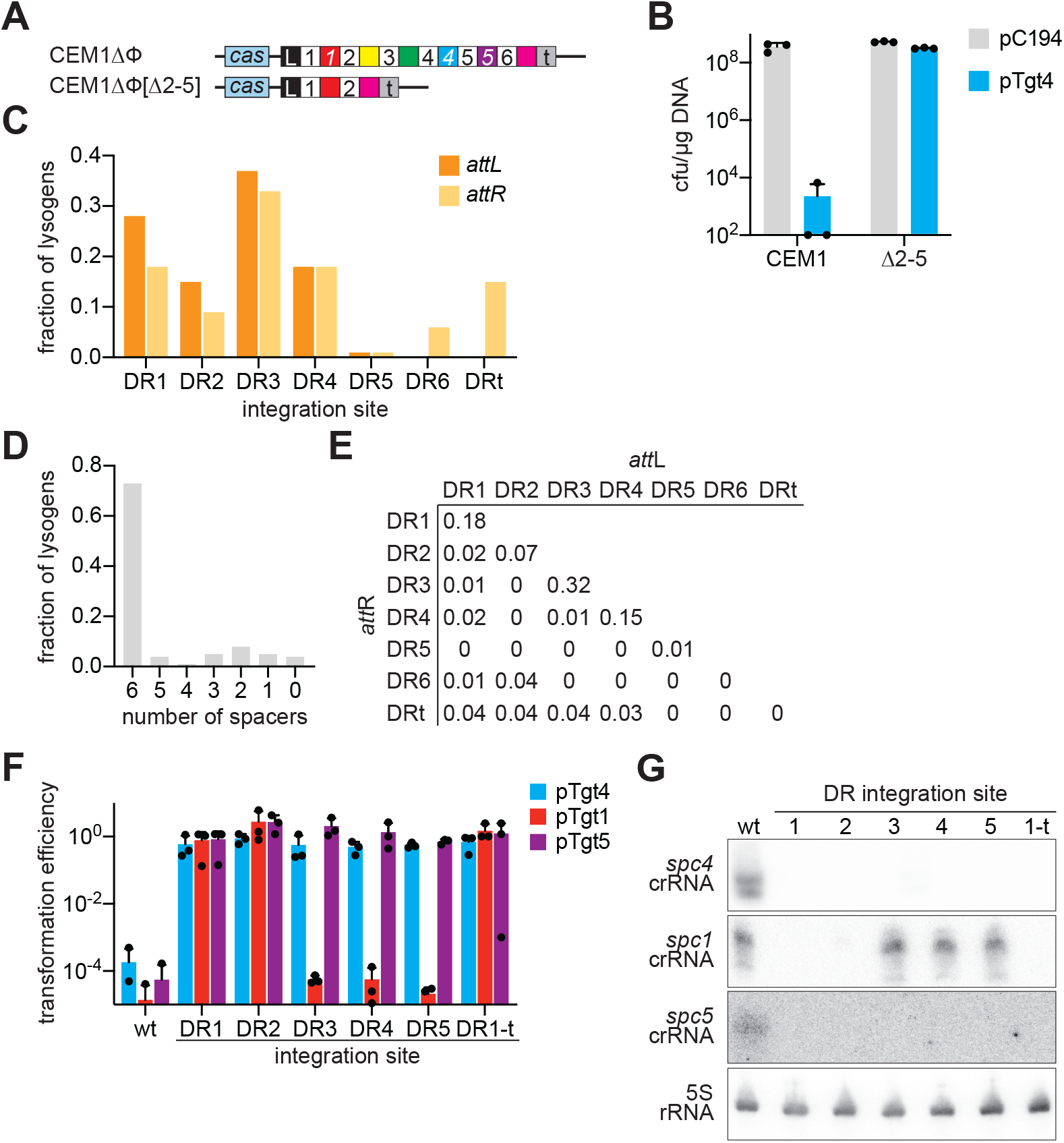
Disruption of CRISPR function by ΦAP1.1 lysogenization. (**A**) *S. pyogenes* CEM1ΔΦ type II-A CRISPR-*cas* locus, wild-type and the mutant lacking spacer-repeat units 2, 3, 4 and 5, [Δ2-5]. (**B**) Transformation efficiency of the pC194 plasmid or a modified version harboring a target for *spc4* from ΦAP1.1, pTgt4, after electroporation of wild-type or [Δ2-5] *S. pyogenes* CEM1ΔΦ competent cells. Mean + STD of 3 biological replicates are reported. (**C**) Frequency of lysogens (n=100) carrying *attL* or *attR* sites in each of the DRs of the CEM1ΔΦ CRISPR locus. (**D**) Frequency of lysogens (n=100) carrying different number of spacers. (**E**) Same as (**C**) but showing the combined distribution of *attL* or *attR* sites. (**F**) Transformation efficiency (relative to a pC194 vector control) of plasmids harboring targets for *spc4*, *spc1* or *spc5* (pTgt4, pTgt1, pTgt5, respectively) after electroporation of wild-type or lysogenic *S. pyogenes* CEM1ΔΦ competent cells. The ΦAP1.1 integration site for each lysogen is indicated. Mean + STD of 3 biological replicates are reported. (**G**) Northern blot analysis of *spc4*, *spc1* and *spc5* crRNAs, as well as 5S rRNA, produced by the strains transformed in (**F**).

We also measured type II-A immunity mediated by *spc1* and *spc5* through transformation of additional plasmids harboring targets for each of these spacers (Fig. 2F). Immunity by *spc1* was abolished by the presence of prophages integrated into DR1 and DR2; i.e., immediately upstream and downstream of the targeting spacer. For *spc5*, all prophages were integrated in upstream DRs and eliminated efficient targeting. Taken together these data suggest that all spacers downstream of the prophage integration site lose their ability to provide immunity, most likely due to interruption of the transcription of the CRISPR array. Also, the spacer immediately upstream of the prophage integration site is ineffective, presumably due to the elimination of repeat sequences that are needed in the pre-crRNA to interact with the tracrRNA during crRNA processing (Deltcheva et al., 2011). To explore this hypothesis, we performed Northern blots to detect *spc4*, *spc1* and *spc5* crRNAs in the different lysogens and found that all lysogens that lacked type II-A immunity in the transformation assay of Figure 2F did not produce the targeting mature crRNA (Fig. 2G). Therefore, we conclude that the integration of ΦAP1.1 in DR1 through DR5 occurs because it prevents the generation of *spc4* crRNA and thus Cas9 cleavage of the residing prophage. Furthermore, we corroborated this pattern of integration by infecting *S. pyogenes* strain NCTC13743 (accession number NZ_LS483384), in which the type II-A CRISPR array contains three spacers that target ΦAP1.1: *spc2*, *spc6* and *spc9* (Figs. S1B, S2A, S2B). We first verified that the type II-A CRISPR system present in this strain is capable of restricting a plasmid containing the three target sequences present in ΦAP1.1 (Fig. S2C). We then infected this triple-targeting strain with ΦAP1.1 and analyzed the prophage location in 25 different lysogenic colonies. We found that a majority of phage integrated into DR2 or DR3; i.e., immediately upstream or downstream of *spc2*, thus neutralizing the immunity provided by this spacer as well as that of the two other downstream targeting spacers (Fig. S2D). Integration events also occurred in repeat sequences downstream of DR3, however they were accompanied by deletions encompassing the upstream targeting spacers. Altogether these data demonstrate that lysogenization of ΦAP1.1 into the repeats of a CRISPR array that harbors targeting spacers neutralizes these spacers, either by preventing the generation of the targeting crRNA or though deletion of the targeting spacer.

### ΦAP1.1 harbors an inhibitor of the *S. pyogenes* type II-A CRISPR-Cas system

Although lysogenization can eventually block targeting after prophage integration, the phage should still be vulnerable to Cas9 recognition and cleavage during DNA injection, (Modell et al., 2017) and even shortly after phage integration, as molecules of Cas9 loaded with *spc4* crRNA will be present in the host cell for a period of time. It is possible a small number of infecting phage stochastically avoid Cas9 cleavage before integration. If this is the case, it will be expected that hosts targeting ΦAP1.1 will display a decrease in the frequency of lysogenization compared to hosts that cannot restrict the phage. To test this, we enumerated lysogens after infection of CEM1ΔΦ and CEM1ΔΦ[Δ2-5] streptococci (Fig. 3A). Surprisingly, we found a minimal effect of type II-A immunity on ΦAP1.1 lysogenization. The same outcome was observed when the triple-targeting *S. pyogenes* strain NTCT13743 and its non-targeting mutant derivative NTCT13743[Δ2-10] were infected with ΦAP1.1 (Fig. S3A). Therefore, both results suggested the presence of a mechanism that avoids Cas9 targeting pre-integration.

**Figure 3.**
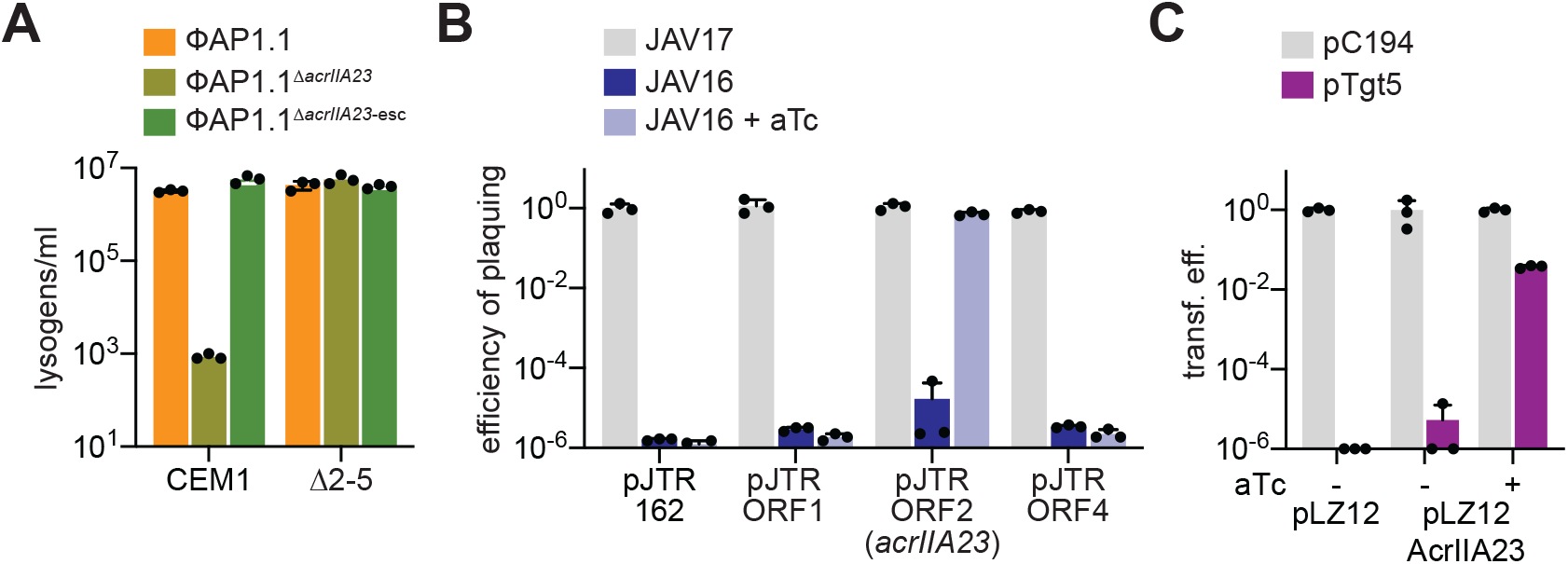
ΦAP1.1 carries a type II-A CRISPR-Cas inhibitor, AcrIIA23. (**A**) Lysogenization rates of ΦAP1.1, ΦAP1.1^*ΔacrIIA23*^, or ΦAP1.1^*ΔacrIIA23*-esc^ after infection of wild-type or [Δ2-5] *S. pyogenes* CEM1ΔΦ cells. Mean + STD of 3 biological replicates are reported. (**B**) Efficiency of plaquing of phage ΦNM4γ4 on soft agar lawns of *S. aureus* JAV17 or JAV16, in the presence or absence of aTc, carrying the pJTR162 control plasmid or versions that express the different ORFs of the ΦAP1.1 anti-CRISPR locus. Mean + STD of 3 biological replicates are reported. (**C**) Transformation efficiency of the pC194 plasmid or a modified version harboring a target for *spc5*, pTgt5, after electroporation of *S. pyogenes* CEM1ΔΦ competent cells carrying the pLZ12 vector or a version expressing AcrIIA23 under the control of a tetracycline-inducible promoter, in the presence or absence of aTc. Mean + STD of 3 biological replicates are reported.

ΦAP1.1 contains a four-gene locus with a similar arrangement to previously identified anti-CRISPR (*acr*) loci (Pawluk et al., 2016; Stanley et al., 2019); i.e., it occurs at the end of the phage genome and it is transcribed in the opposite direction of the lytic genes (Fig. S1B). Due to difficulty of detecting plaque-forming units (pfus) after treatment of *S. pyogenes* with ΦAP1.1, we decided to determine if any of these four open reading frames (ORFs) inhibit the type II-A CRISPR-Cas response against phage infection in a *Staphylococcus aureus* heterologous system. To do this we inserted the *S. pyogenes* CEM1ΔΦ type II-A system into the lipase gene of *S. aureus* RNA4220 using the integrative vector pCL55 (Lee et al., 1991), generating strain JAV17. Additionally, we inserted a CRISPR locus with a spacer matching the *gp68* gene from the staphylococcal phage ΦNM4γ4 (Varble et al., 2019), generating JAV16 (Fig. S3B). JAV17 and JAV16 strains were transformed with each of the individual ΦAP1.1 ORFs under the control of an anhydrotetracycline (aTc)-inducible promoter, on the pJTR162 staphylococcal plasmid backbone (Rostol and Marraffini, 2019), with the exception of *ORF3*, for which we did not obtain transformants. Strains harboring the pJTR-ORF1, pJTR-ORF2 and pJTR-ORF4 constructs were then infected with ΦNM4γ4 in the presence or absence of aTc, and pfus were counted (Fig. 3B). We found that none of the ORFs significantly affected the immunity of the JAV16 strain in the absence of the inducer, and ΦNM4γ4 propagation was limited. In the presence of aTc, however, *ORF2* expression enabled high levels of viral replication, similar to those observed in the absence of type II-A CRISPR-Cas immunity after infection of the JAV17 strain. ORF2 did not display amino-acid homology to previously identified Acrs, and therefore we named it AcrIIA23, according to the current nomenclature (Bondy-Denomy et al., 2018). To corroborate that AcrIIA23 is active in *S. pyogenes*, we transformed strain CEM1ΔΦ with the streptococcal plasmid pLZ12 (Perez-Casal et al., 1991) or a version containing AcrIIA23 under the control of an aTc inducible promoter, pLZ12-AcrIIA23. To measure type II-A immunity, each of these strains was then transformed with a second plasmid containing the *spc5* target sequence, pTgt5, or an empty vector control, pC194, in the presence or absence of aTc (Fig. 3C). We found that in the absence of AcrIIA23 expression there was low transformation efficiency for pTgt5 and that this was reverted to the high levels observed for the non-target pC194 control by the addition of aTc. Altogether these results demonstrate that ΦAP1.1 contains a novel inhibitor of the *S. pyogenes* CRISPR-Cas immune response.

### AcrIIA23 is required for ΦAP1.1 lysogenization under targeting conditions

To investigate the importance of AcrIIA23 for ΦAP1.1 lysogenization under targeting conditions, we generated a phage with a deletion of the inhibitor gene, ΦAP1.1^Δ*acr23*^. We then infected strains CEM1ΔΦ and CEM1ΔΦ[Δ2-5] with the mutant virus and measured the formation of lysogens (Fig. 3B). We observed that, as opposed to the wild-type ΦAP1.1, integration into the CRISPR array of strain CEM1ΔΦ, but not in that of CEM1ΔΦ[Δ2-5], was markedly impaired. A similar decrease in lysogenization was detected when strain NTCT13743 was infected with the ΦAP1.1^Δ*acr23*^ phage (Fig. S3A). These results suggest that Cas9 cleavage of the phage genome in the absence of the inhibitor prevents lysogenization. To corroborate this, we isolated a ΦAP1.1^Δ*acr23*^ virus that was able to escape targeting by *S. pyogenes* CEM1ΔΦ bacteria, which contained a mutation in the PAM sequence of the *spc4* target (Fig. S1D) to escape Cas9 cleavage (Jinek et al., 2012; Nussenzweig et al., 2019). We then measured the lysogenization frequency of ΦAP1.1^Δ*acr23*-esc^ and also found equally high levels of lysogen formation in the absence of the inhibitor (Fig. 3A). Importantly, as opposed to wild-type ΦAP1.1, this mutant phage was capable of integrating its *attL* into DR6 and DRt (Fig. S4A, B) to produce lysogens in which the targeting *spc4* crRNA is generated (Fig. S4C), providing efficient restriction of pTgt4 (Fig. S4D). Based on these results, we conclude that, through the inhibition of type II-A CRISPR-Cas targeting, AcrIIA23 protects the phage during the early stages of infection, allowing ΦAP1.1 integration into the CRISPR array.

### *acrIIA23* transcription is turned off during ΦAP1.1 lysogenization

In spite of the presence of an active inhibitor of the type II-A CRISPR-Cas immune response, the ΦAP1.1 prophage does not prevent plasmid targeting if the crRNA is still produced by the host (Fig. 2F). In addition, integration only occurs when immunity is neutralized by the deletion of the targeting spacers (Fig. S2D) or by the elimination of the crRNA guides they produce (Fig. 2G). In other words, AcrIIA23 activity is not used to prevent “auto-immunity” against the prophage. This is a striking contrast to other Acrs found in prophages, which are expressed and active during both the lytic and lysogenic stages of infection (Bondy-Denomy et al., 2013; Stanley et al., 2019). We performed next-generation RNA sequencing (RNA-seq) to determine the pattern of expression of *acrIIA23*. During the lytic cycle, we detected expression of a transcript spanning ORF1 and the inhibitor as early as 5 minutes after infection, continuing for at least 90 minutes and resulting in a constant increase of the mRNA levels (Fig. 4A). The same pattern is observed for a second transcriptional unit containing ORF3 and ORF4. In contrast, when we performed RNA-seq of a lysogen where ΦAP1.1 was integrated into DR3, we found that only the ORF3-ORF4 mRNA is produced, while both the ORF1 and *acrIIA23* genes are silenced (Fig. 4B). These results demonstrate that ΦAP1.1 harbors a conditional inhibitor that is transcribed, and thus active, during the lytic but not during the lysogenic cycles of this temperate phage.

**Figure 4.**
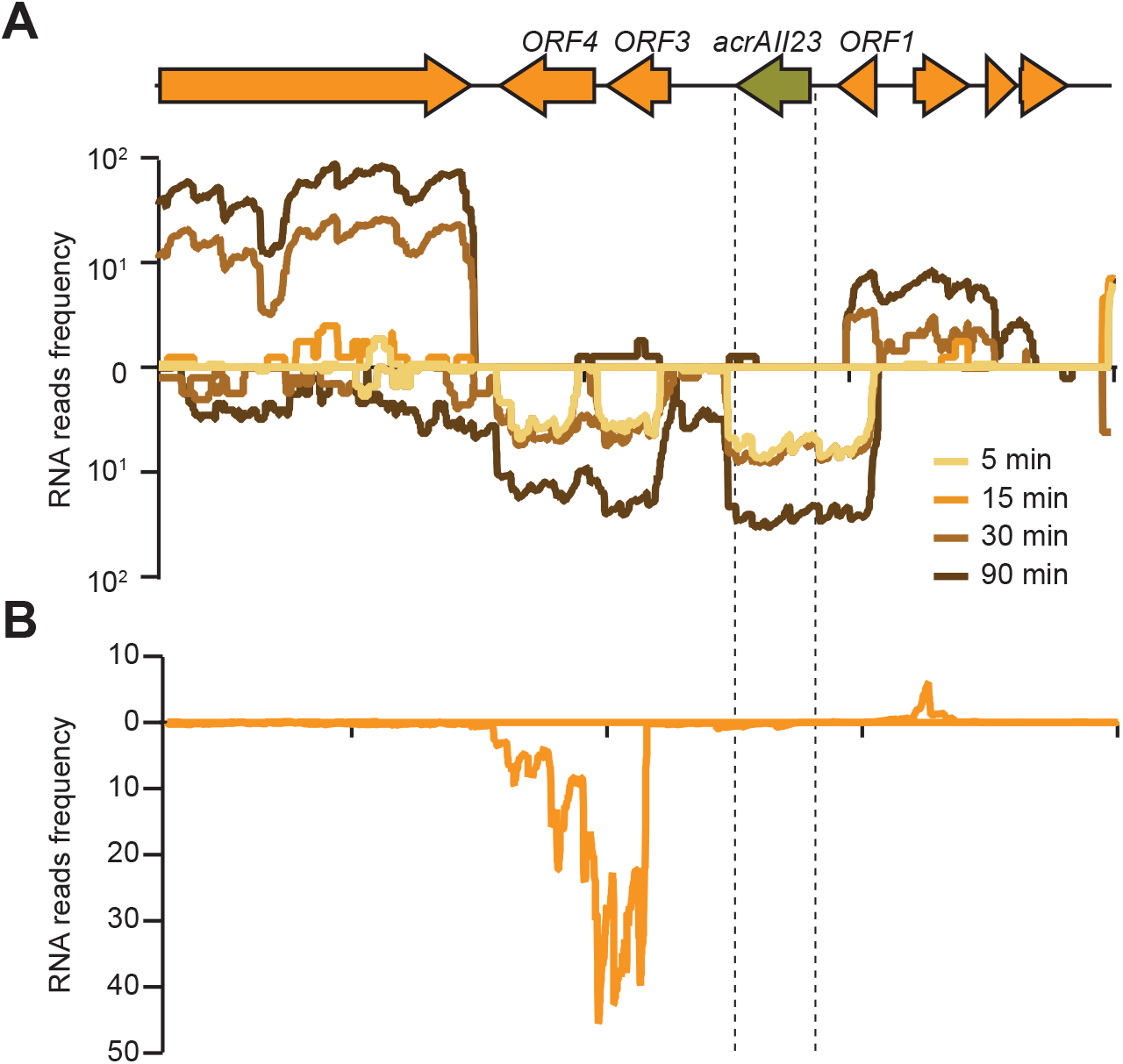
AcrIIA23 is not transcribed from ΦAP1.1 prophages. (**A**) RNAseq of *S. pyogenes* CEM1ΔΦ cells at different times after infection with ΦAP1.1. The fraction of reads (relative to the total number of reads) that map to the *acr* region of the phage are plotted. (**B**) Same as (**A**) but using RNA from a culture of *S. pyogenes* CEM1ΔΦ (ΦAP1.1::DR3) lysogen.

### Aberrant ΦAP1.1 excision during prophage induction can transduce immunity

In most of the strains sequenced to date, ΦAP1.1-like phages are located in a spacer-less CRISPR locus, flanked by a partial DR and a DR’ (Fig. S1A). In our lysogenization experiments we observed that integration into a full CRISPR array can lead to spacer deletions (Figs. 1E, 2D, S2D). We hypothesized that the cycling between integration and excision could eventually lead to the elimination of all spacers and the prophage insertion pattern typically observed in sequenced genomes. To investigate this, we grew two CEM1ΔΦ (ΦAP1.1) lysogens, with prophage insertions in DR3 and DR5, for 24 hours in liquid media containing mitomycin C (MMC) to trigger rounds of induction and integration. We then collected bacteria and amplified the *attL* and *attR* sites to detect changes in the number of spacers that flank each integration site. For the prophages originally inserted into DR3, we did not detect changes in the *attL* site, but the *attR* site appeared in PCR products of smaller sizes that indicated a location more proximal to the terminal repeat, either due to the deletion of spacer-repeat units, or re-insertion into repeats downstream of DR3 (Fig. 5A). Importantly, these new PCR products were diminished in the absence of MMC treatment, indicating that they did not arise from pre-existing repeat-spacer deletions present in the culture, and are not an artifact of amplification. A similar result was obtained with the prophage originally inserted in DR5, for which we were able to detect a smaller product, whose size indicated integration of the *attR* site into the DR’ (Fig. 5B). Moreover, amplification of the *attP* site of the phages induced with MMC produced a DNA fragment consistent with the size of an additional spacer-repeat unit, which was not seen when DNA from the ΦAP1.1 stock was used as template (Fig. 5C). These results suggest that over evolutionary timescales multiple rounds of prophage integration and excision can lead to the eventual erasure of the CRISPR array observed in the natural isolates of ΦAP1.1 lysogens.

**Figure 5.**
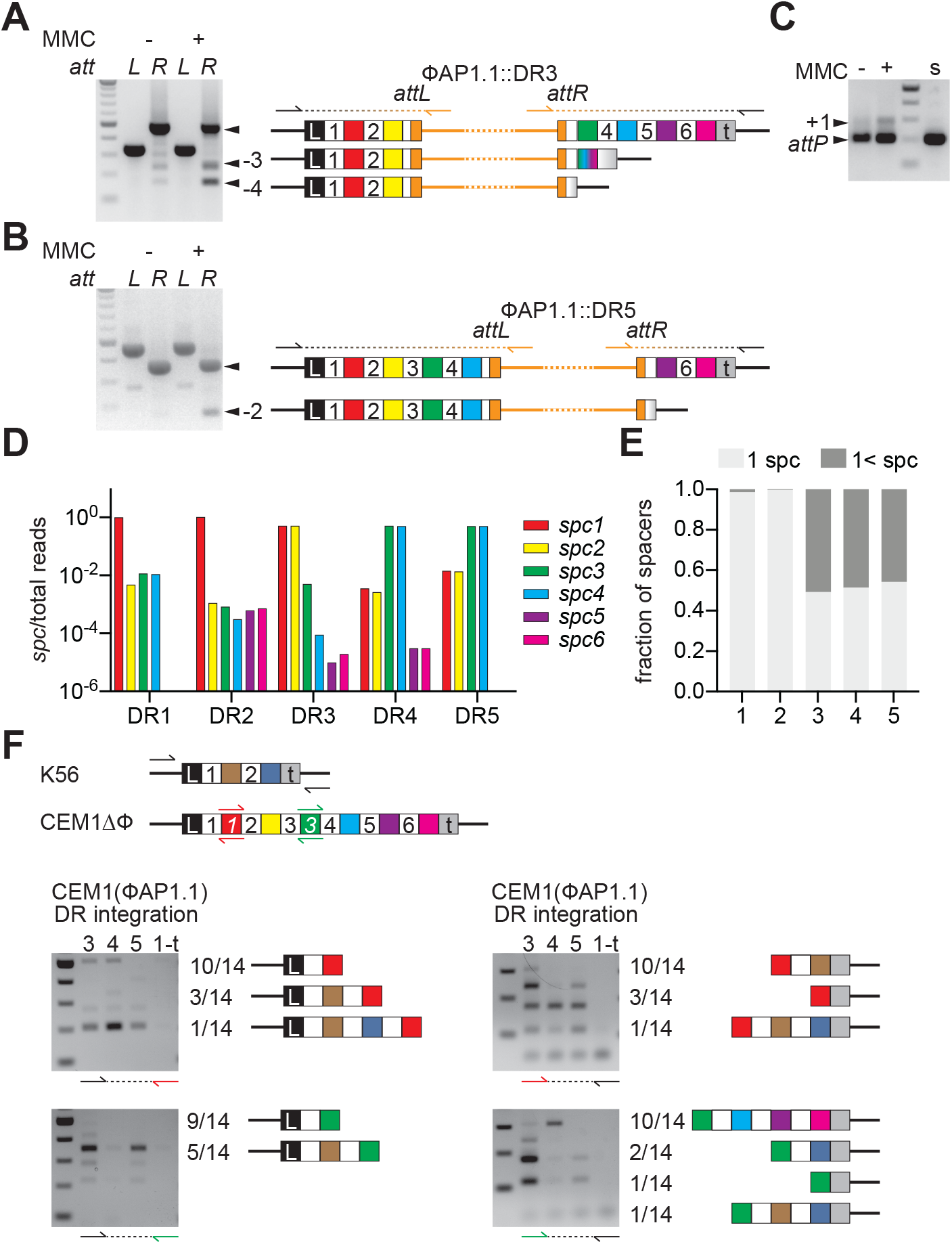
ΦAP1.1 mediates the transduction of CRISPR spacers. (**A**) Agarose gel electrophoresis of the *attL* or *attR* PCR products obtained after treatment of *S. pyogenes* CEM1ΔΦ (ΦAP1.1::DR3) lysogens with mitomycin C (MMC). Primers used and the expected products are shown in the diagrams to the right. Black arrowheads indicate PCR products that lack a number of the expected spacer-repeat units. (**B**) Same as (**A**) but after treatment of *S. pyogenes* CEM1ΔΦ (ΦAP1.1::DR5) lysogens. (**C**) Same as (**A**) but after amplification of the ΦAP1.1 *attP* site. “s” indicates a control amplification of a phage stock that has never been lysogenized. (**D**) DNAseq of the *attP* amplification products obtained via PCR after induction of ΦAP1.1 prophages integrated in the indicated DR. The fraction of reads (relative to the total reads) corresponding to each spacer sequence associated with the *attP* site is shown. (**E**) Distribution of the number of spacers associated with the *attP* in (**D**). (**F**) Agarose gel electrophoresis of PCR products obtained from the amplification of the CRISPR locus of *S. pyogenes* K56 lysogens generated after infection with lysates obtained from *S. pyogenes* CEM1ΔΦ lysogens where ΦAP1.1 was integrated into DR3, DR4, DR5 or DR1-t (lacking all spacers). Top diagram shows the K56-, *spc1*- or *spc3*-specific primers used as black, red or green arrows, respectively. Diagrams next to gels show the spacer composition of 14 different PCR products that were sequenced, obtained after transduction of *S. pyogenes* CEM1ΔΦ (ΦAP1.1::DR3) lysates.

The possibility that spacers are deleted during ΦAP1.1 induction, raises the question of whether these spacers can become part of the viral genome during prophage excision. In principle, every DR and DR’ contains both *attR* and *attL* sites (Figs. 1A) and therefore can be used by the phage integrase during excision. To determine if spacers can be excised along with the viral genome, CEM1ΔΦ (ΦAP1.1) lysogens with the prophage integrated into DR3 (Figs. S5A) or DR5 (Figs. S5B) were induced with MMC, viral particles were purified from culture supernatants, and their DNA extracted to be used as template for *spc1*-(Fig. S5C) or *spc2*-specific (Fig. S5D) PCR assays. In all cases we found PCR products that indicated the presence of spacer-repeat units in the DNA of the viral particles, with the intensity of the amplicons indicating more *spc1*-than *spc2*-containing viral particles. This suggested a possible preference for certain excision products. To obtain a more accurate representation of the population of spacer-carrying phages, we amplified the *attP* from the DNA of the purified phage particles collected after the MMC induction of prophages inserted into DR1-5. While the most prominent product consisted of the expected *attP* without any additional spacer-repeat units (Fig. 5C), we purified the DNA corresponding to larger amplicon sizes and subjected them to next generation sequencing to obtain the relative fraction of reads of each spacer (Fig. 5D). We observed a “proximity” effect, whereby the spacer sequences most commonly found in excised viral genomes were the closest to the DR in which the prophage was originally integrated; i.e., *spc1* was the most commonly acquired spacer by phages previously residing in DR1 and DR2, *spc1* and *spc2* for DR3, and *spc3* and *spc4* for DR4 and DR5. This trend, however, was not perfectly uniform and therefore the results suggest that there are certain hotspots where *attP* formation is preferred. This was also reflected in the number of repeat-spacer units that were excised along with the prophage. Whereas prophages integrated into DR1 and DR2 typically contained a single unit, at least half of the reads of the DR3-5 lysogens contained two or more spacers (Fig. 5E).

The presence of repeat-spacer units in viral particles suggested that these could be added to the CRISPR array of new hosts following lysogenization (Fig. S5E). To test this, we induced with MMC CEM1ΔΦ (ΦAP1.1) lysogens in which the prophage was integrated into DR3, DR4, DR5 or a non-lysogen as a control. Viral particles were filtered from the culture supernatants and used to infect strain K56. Lysogens that survived superinfection were collected after five hours and their genomic DNA was extracted to use as template for spacer-specific PCR assays. We used primers that anneal outside of K56 CRISPR array, in the 5’ or 3’ flanking regions, in combination with primers that anneal to either the top or bottom strand of *spc1* or *spc3* sequences from the CEM1ΔΦ CRISPR locus (Fig. S5E). Amplification resulted in many different products for each lysogen population with different numbers and combinations of repeat-spacer units (Fig. 5F). This was most likely a consequence of the presence of several repeat sequences in both phage and bacterial genomes, each being a suitable *attP* and *attB* site, respectively, thus multiplying the possibilities for integration (Fig. S5E). From this data we conclude that all of the three ΦAP1.1 populations were able to transduce *spc1* and *spc3* from the CEM1ΔΦ strain into the CRISPR locus of the K56 strain. Sanger sequencing of PCR products obtained after transduction with a ΦAP1.1 prophage previously integrated in the DR3 of CEM1ΔΦ, confirmed this conclusion (Fig. 5F).

## Discussion

Here we describe the interaction between the temperate phage ΦAP1.1 and the type II-A CRISPR-Cas system of its host, *S. pyogenes*. Uniquely, this phage uses the CRISPR DR sequence as the attachment site for integration and expresses an anti-CRISPR, AcrIIA23, that it is only active during the lytic portion of its replication cycle. This unique mode of insertion generates a very complex and rich molecular interplay between the virus and CRISPR immunity. Integration disrupts the transcription of the crRNAs located downstream of the prophage and eliminates the portion of the repeat sequence important for crRNA maturation of the spacer immediately upstream of the *attL*. However, the transcription of spacers further upstream and of the *cas* genes (in all *S. pyogenes* strains sequenced to date the *cas* operon is located upstream of the type II-A CRISPR array) is not altered. As a consequence of this, ΦAP1.1 integration can selectively turn off the immunity provided by different sets of spacers (Fig. 2F). This differs from the mode of action of previously described anti-CRISPRs that broadly turn off CRISPR function (Davidson et al., 2020), and can provide a selective advantage for ΦAP1.1 (and related phages), which could exploit integration to neutralize spacers that target its genome, but at the same time maintain the immunity provided by other spacers against competitor phages, promoting host survival. Additionally, these lysogens could also retain the ability to acquire new spacers in the first position of the CRISPR array against new invaders. Furthermore, due to the presence of multiple *attL* and *attR* sites, one in every upstream and downstream DR, respectively, induction of the ΦAP1.1 prophage can result in the incorporation of repeat-spacer units into the viral genome. We observed that different spacer-repeat units were excised along ΦAP1.1 genome at different levels, and therefore we speculate, as it is the case for the *att* sites of other temperate phages, that the excision of the *attR* and *attL* elements is influenced by its flanking sequences, which would be the CRISPR spacers in the case of ΦAP1.1. Once in the viral genome, repeat-spacer units have the possibility to be transduced into a different *S. pyogenes* strain and, depending on their position relative to the prophage, provide new immunity to the recipient cell (Fig. 5F). Such spacer transduction events would increase the range of viral targets in the CRISPR system in an adaptation-independent manner, and benefit both the residing prophage and its host. In contrast, we and others have previously reported transduction of entire CRISPR systems (Varble et al., 2019; Watson et al., 2018), which is enhanced by recombination between targeting spacers in the CRISPR array and the phage genome. While this phenomenon is due to the promiscuity of phage recombinases, ΦAP1.1-mediated transduction of spacers appears to be a specifically evolved interaction that intertwines the phage life-cycle with the CRISPR locus.

*S. pyogenes* prophages carry virulence genes that play fundamental roles in the different manifestations of group A streptococcus (GAS) disease (Banks et al., 2002). Both the gain- and loss-of-function of spacers that result from integration of ΦAP1.1 into the CRISPR array could play an important role in the restriction or tolerance of prophages targeted by type II-A immunity. Therefore, we believe that the diversity of spacer genotypes generated during the interplay between ΦAP1.1 phages and their hosts, even if occurring at a very low frequency, could lead to the selection of prophage-dependent phenotypes that enable specific *S. pyogenes* adaptations, including the modulation of its virulence. However, in most of the *S. pyogenes* strains sequenced so far, ΦAP1.1-related prophages are integrated into a single DR’ element (Fig. S1A). We suggest that after the initial insertion into a full CRISPR array, subsequent rounds of excision and integration lead to the selection of spacer deletions that ultimately erase all repeat-spacer units. This would cause an irreversible inhibition of the type II-A CRISPR-Cas immune response, since the single DRt that can be re-formed after prophage excision does not support spacer acquisition (data not shown). Thus, it seems that from all the possible ΦAP1.1 lysogens that can be formed, natural selection has favored those that lack type II-A anti-phage immunity. We hypothesize that this decreases barriers against viral infection, facilitating the incorporation of the multiple prophages, and therefore allowing lateral gene flow that plays a critical role for *S. pyogenes* survival and colonization of the human host.

While ΦAP1.1 integration can neutralize CRISPR immunity, inhibition only occurs after lysogenization. Therefore, the phage could be susceptible to Cas9 cleavage during genome injection if there is a targeting spacer in the type II-A CRISPR array. To prevent this, ΦAP1.1 expresses an inhibitor, AcrIIA23, immediately after infection. Transcription of the anti-CRISPR, however, is completely turned off after prophage integration. This is in contrast to other Cas9 inhibitors, which remain active in the lysogen. Indeed, the first *acrIIA* genes were discovered after the identification of type II-A CRISPR systems harboring seemingly inactive spacers with perfect targets in residing prophages of *Listeria* (Rauch et al., 2017). This is also the case for *acrI* genes carried by *Pseudomonas* prophages, which are able to inhibit the type I CRISPR response (Bondy-Denomy et al., 2013). Following robust transcription at the onset of phage infection, *acr* genes are repressed by anti-CRISPR associated (Aca) transcription factors to prevent the deleterious effects of the *acr* high transcription on the downstream phage genes (Osuna et al., 2020; Stanley et al., 2019). The acr region of ΦAP1.1 contains two genes, *orf3* and *orf4*, which are transcribed during lysogeny. Since *aca* genes are invariably found in the vicinity of *acr* genes (Marino et al., 2018; Stanley et al., 2019; Yin et al., 2019), we hypothesize that *orf3* or *orf4* produce the Aca that regulates transcription of *acrIIA23*, providing a tight repression that is not observed in other lysogens that produce basal levels of inhibitor. Regardless of the mechanism that prevents *acrIIA23* transcription in ΦAP1.1 lysogens, the absence of Cas9 inhibition after integration only selects prophage insertions that prevent the generation of the targeting crRNA. In summary, our study presents a unique dynamic in the parasite-host conflict, where the phage can modulate the CRISPR immunity of streptococci to protect itself from targeting, while still allowing the defense of the lysogen against other invaders.

## Acknowledgements

We thank Pascal Maguin and Alexander J Meeske for bioinformatic assistance with the RNA-seq analysis. Support for this work comes from the National Institute of Health Director’s Pioneer Award 1DP1GM128184-01 and the Burroughs Wellcome Fund PATH Award to LAM. LAM is an Investigator of the Howard Hughes Medical Institute. AV was supported by the Arnold and Mabel Beckman Postdocotoral Fellowship. JTR is supported by the Boehringer Ingelheim Fonds PhD fellowship.

## Author contributions

AV and LAM conceived the study. EC, CWE, and VAF aided with experimental design. AV performed all the experiments with the help of JF. EC assisted with bioinformatic analysis and EC and CWE provided assistance, strains and reagents for *S. pyogenes* experiments. JTR provided reagents. AV and LAM wrote the manuscript with the help of the other authors.

---

**Figure S1.**
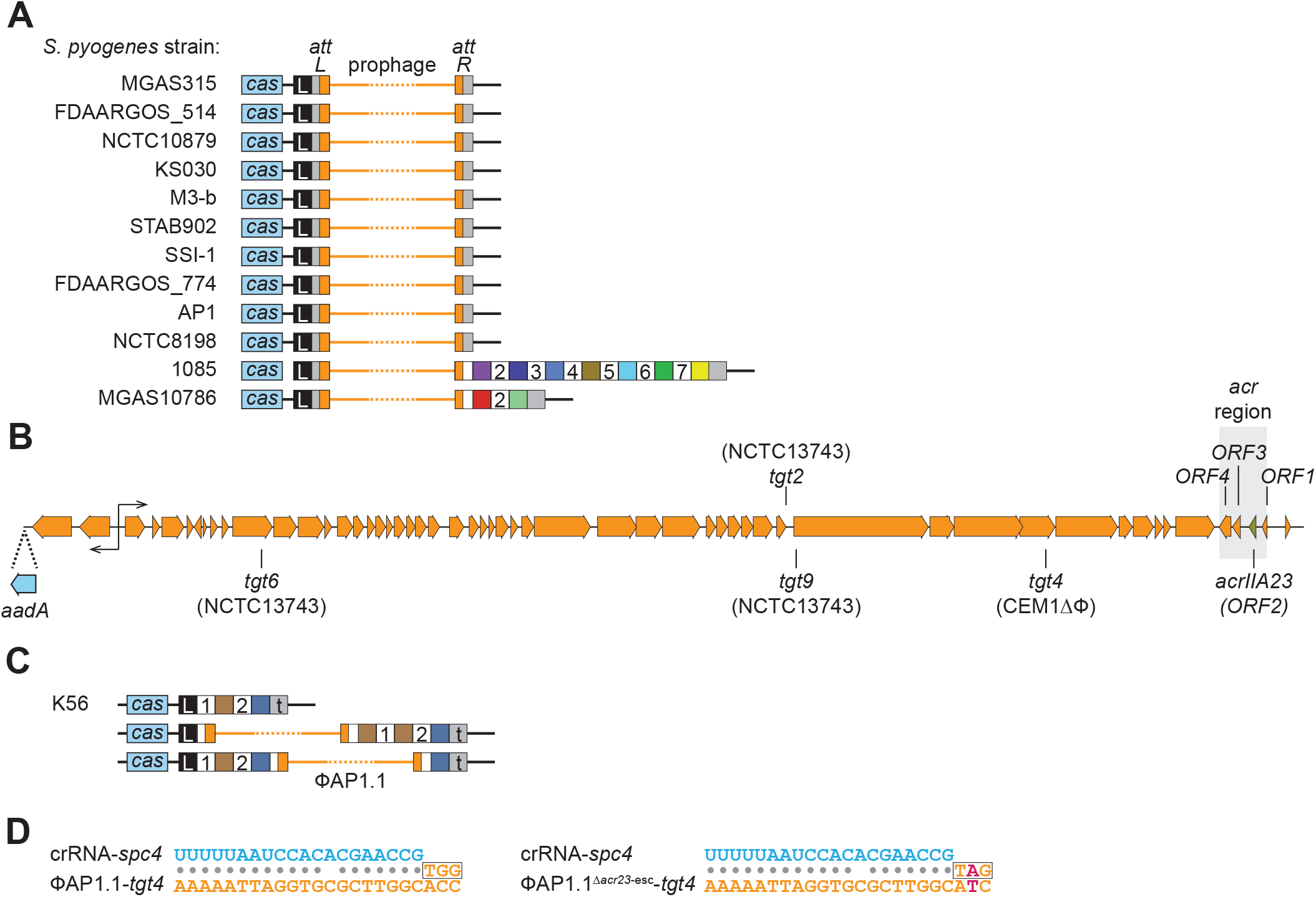
ΦAP1.1 integration into *S. pyogenes* type II-A CRISPR loci. (**A**) ΦAP1.1-like phages integrated into the type II-A CRISPR locus of *S. pyogenes* strains. (**B**) Schematic of the ΦAP1.1 genome. Targets for the spacers found in different strains are shown on top or bottom depending on the phage DNA strand that is targeted. The *acr* region and its four ORFs are highlighted in grey. The *aadA* gene inserted to confer spectinomycin resistance to ΦAP1.1 lysogens is shown in light blue. (**C**) Spacer duplication events observed in *S. pyogenes* K56(ΦAP1.1) lysogens. (**D**) Sequences of the *S. pyogenes* CEM1ΔΦ *spc4* crRNA annealed to its targets in ΦAP1.1 and ΦAP1.1^*ΔacrIIA23*-esc^. The PAM nucleotides are boxed. The escape mutation of ΦAP1.1^*ΔacrIIA23*-esc^ is in red.

**Figure S2.**
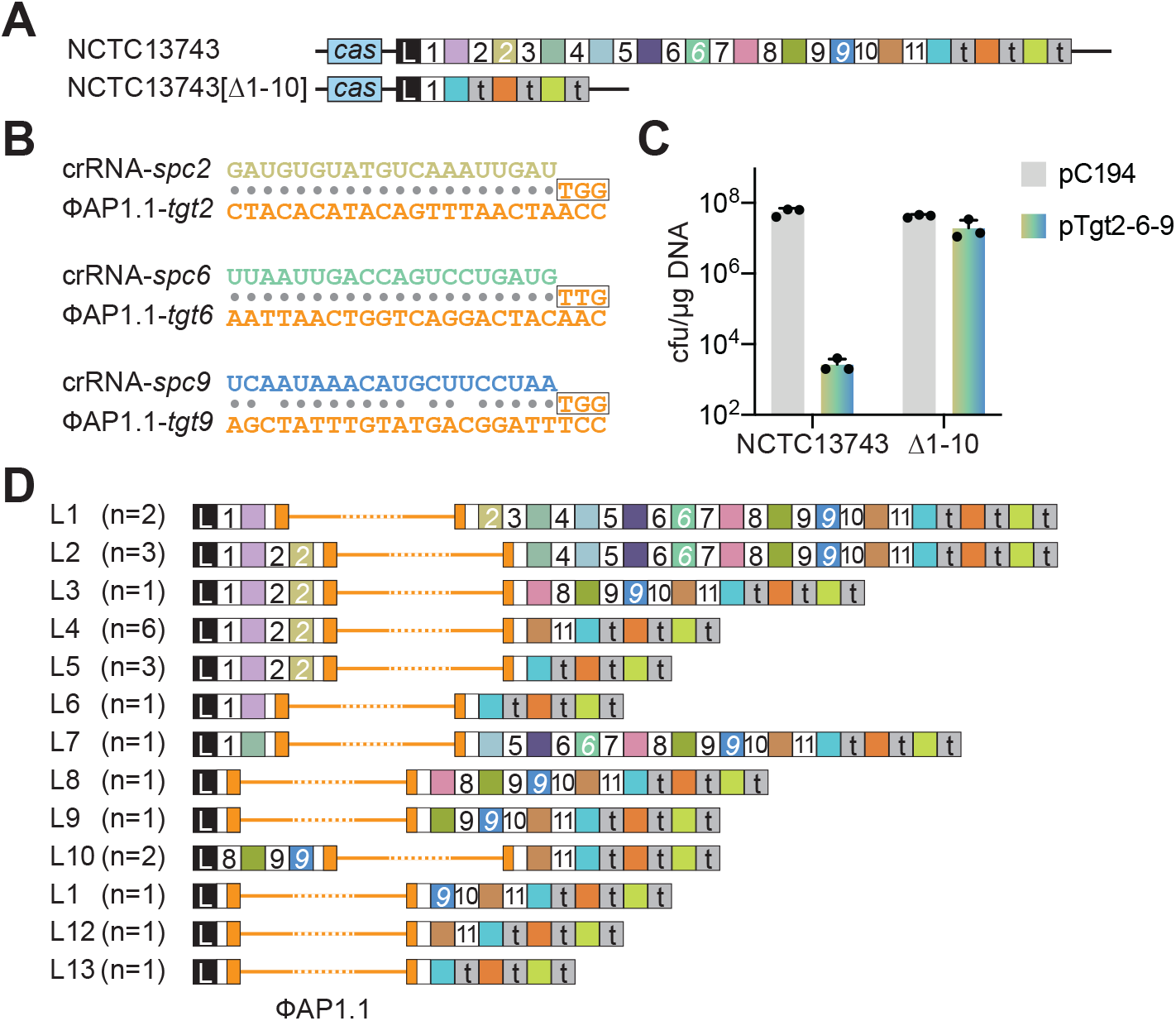
ΦAP1.1 lysogenization into the *S. pyogenes* NCTC13743 type II-A CRISPR locus. (**A**) *S. pyogenes* NTCT13743 type II-A CRISPR-*cas* locus, wild-type and the mutant lacking spacer-repeat units 1 through 10, [Δ1-10]. (**B**) Sequences of the *S. pyogenes* NTCT13743 *spc2*, *spc6* and *spc9* crRNAs annealed to their targets in ΦAP1.1. The PAM nucleotides are boxed. (**C**) Transformation efficiency of the pC194 plasmid or a modified version harboring the three targets for *spc2*, *spc6* and *spc9*, pTgt2-6-9 from ΦAP1.1, after electroporation of wild-type or [Δ1-10] *S. pyogenes* NCTC13743 competent cells. Mean + STD of 3 biological replicates are reported. (**D**) Diagrams of the type II-A locus of 25 ΦAP1.1 lysogens in *S. pyogenes* NCTC13743, reconstructed after sequencing of their *attL* and *attR* sites.

**Figure S3.**
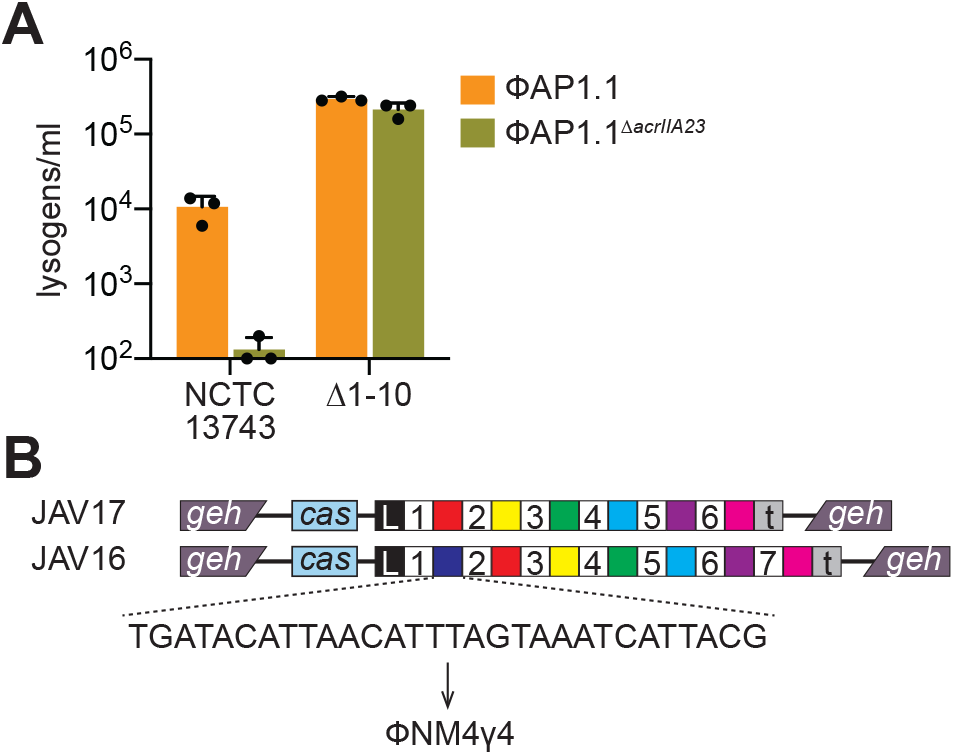
AcrIIA23 inhibits the type II-A CRISPR-Cas response in *S. pyogenes* NCTC13743 and *S. aureus* RN4220. (**A**) Lysogenization rates of ΦAP1.1 or ΦAP1.1^*ΔacrIIA23*^ after infection of wild-type or [Δ1-10] *S. pyogenes* NCTC13743 cells. Mean + STD of 3 biological replicates are reported. (**B**) Integration of the type II-A CRISPR locus of *S. pyogenes* into the lipase gene (*geh*) of *S. aureus* RN4220, generating strain JAV17. Integration of a locus containing a spacer targeting the staphylococcal phage ΦNM4γ4 generates strain JAV16.

**Figure S4.**
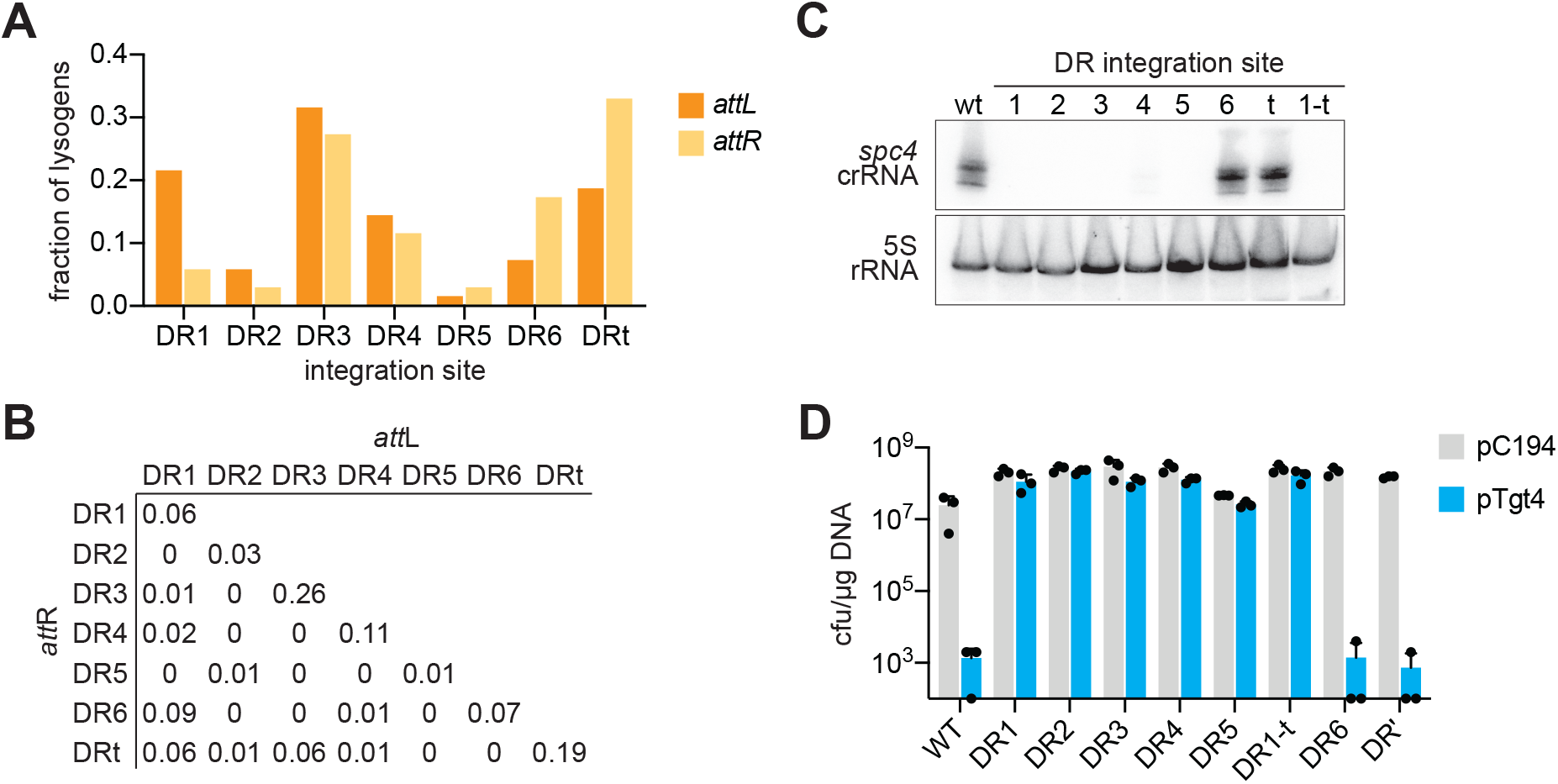
ΦAP1.1^*ΔacrIIA23*-esc^ integration pattern. (**A**) Frequency of ΦAP1.1^*ΔacrIIA23*-esc^ lysogens (n=50) carrying *attL* or *attR* sites in each of the DRs of the CEM1ΔΦ CRISPR locus. (**B**) Same as (**A**) but showing the combined distribution of *attL* or *attR* sites. (**C**) Northern blot analysis of *spc4* crRNAs, as well as 5S rRNA, produced by wild-type *S. pyogenes* CEM1ΔΦ or its different ΦAP1.1 (DR1-5, 1-t) or ΦAP1.1^*ΔacrIIA23*-esc^ (DR6, DRt) lysogens. (**D**) Transformation efficiency of the pC194 plasmid or a modified version harboring a target for *spc4* from ΦAP1.1, pTgt4, after electroporation of the strains used in (**C**). Mean + STD of 3 biological replicates are reported.

**Figure S5.**
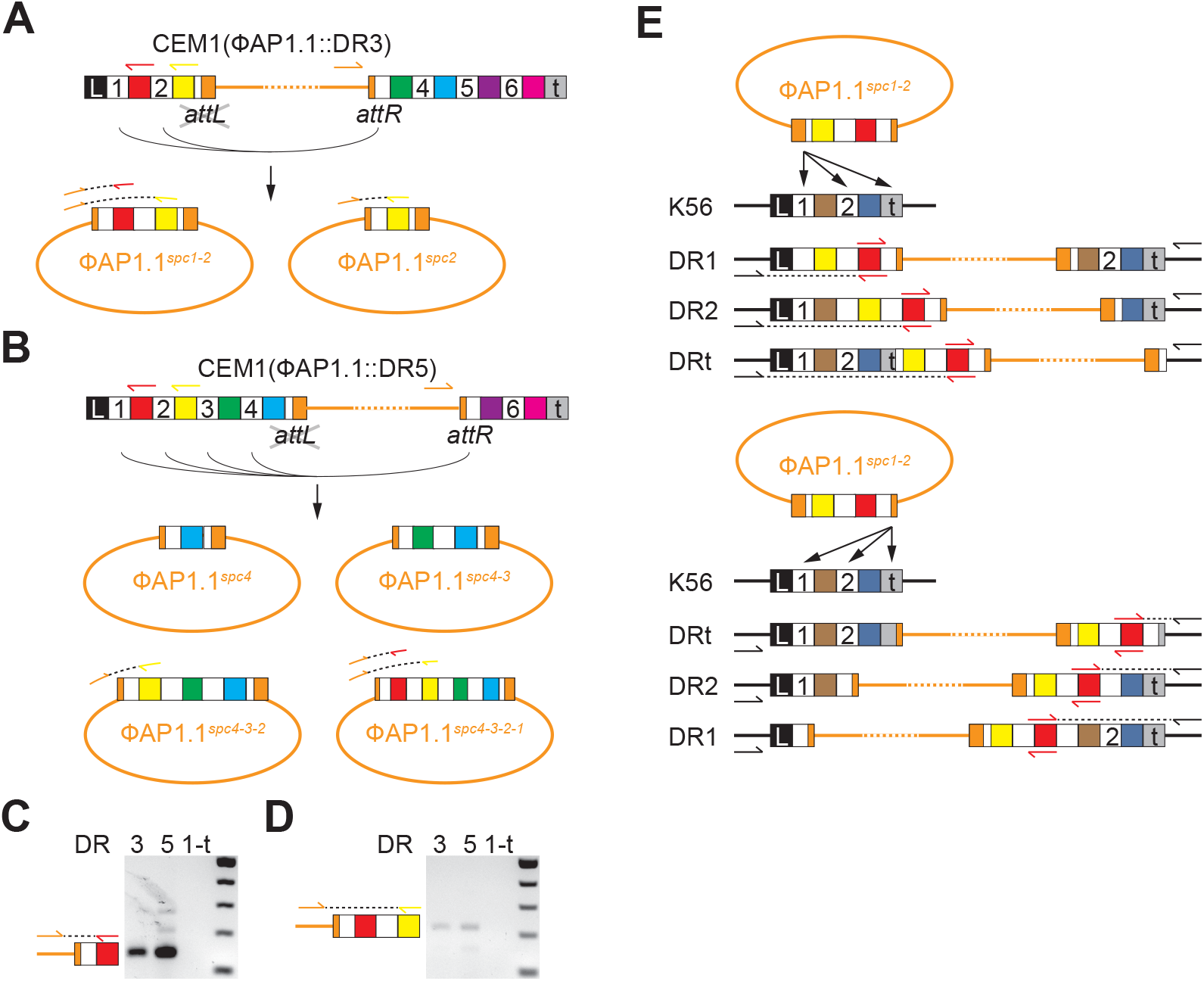
ΦAP1.1-mediated transduction of spacers from the CEM1ΔΦ to the K56 CRISPR locus. (**A**) Example of incorrect ΦAP1.1 excision in *S. pyogenes* CEM1ΔΦ (ΦAP1.1::DR3) lysogens, where DR1 or DR2 are used instead of *attL* for recombination with *attR*. ΦAP1.1-, *spc1*- and *spc2*-specific primers used to detect expanded *attP* sites are shown as orange, red and yellow arrows, respectively. (**B**) Same as (**A**) but for a ΦAP1.1::DR5 lysogen. (**C**) Agarose gel electrophoresis of PCR products obtained after the amplification of the *attP* site of ΦAP1.1 phages induced with MMC from ΦAP1.1::DR3, ΦAP1.1::DR5 and ΦAP1.1::DR1-t lysogens, using *spc1*-specific primers. (**D**) Same as (**C**) but using *spc3*-specific primers. (**E**) Example of possible recombination of the 5’ end (top) or 3’ end (bottom) *attP* site of the ΦAP1.1^*spc1-2*^ phage with the three *attB* sites present in the DR1, DR2 and DRt of *S. pyogenes* K56.

## Methods

### Bacterial strains and growth conditions

Culture of *S. pyogenes* and *S. aureus* was carried out in brain-heart infusion (BHI) medium at 37°C. For *S. pyogenes*, liquid experiments were carried out in 10 milliliters of medium in 15 ml conical tubes unless otherwise noted. *S. pyogenes* media was supplemented with 5 μg/ml chloramphenicol or 50 μg/ml spectinomycin for plasmid maintenance and/or lysogen selection. For *S. aureus*, liquid experiments were carried out in 3 milliliters of medium in 15 ml conical tubes. *S. aureus* media was supplemented with 5 μg/ml chloramphenicol or 10 μg/ml erythromycin for plasmid maintenance and/or strain selection. All strains used in this study are listed in Table S1.

### Induction of Lysogens

To isolate phage, overnight cultures of the indicated lysogens were diluted 1:50 into 10 mls of BHI and grown at 37 °C without agitation. At OD_600_ = 0.3 cells were induced with 0.5 μg/ml of mitomycin C. Four hours post-induction cells were spun down and supernatant was filtered through 0.45-μm syringe filters (Acrodisc).

### Strain construction

To make *S. aureus* strains with the integrated SF370 CRISPR locus (JAV17 and JAV16, Table S1), pCL55-integration plasmids were constructed containing the SF370 *cas* genes with the wildtype CRISPR array (pAV121) or with a CRISPR array containing a ΦNM4γ4-targeting spacer (pAV122) (5’-TGATACATTAACATTTAGTAAATCATTACG −3’). These vectors were transformed in RN4220, recovered at 30°C for 2 hours, and plated on 5 μg/ml chloramphenicol. Integration was confirmed using primers oGG50 and oGG96.

Deletions of spacers in *S. pyogenes* strains CEM and NCTC 13743 were isolated by curing lysogens ΦAP1.1-spec. Prophages were cured by transforming lysogens as described in the transformation assay section with a plasmid containing the SF370 type II-A *cas9* with a CRISPR array targeting the ΦAP1.1-spec, pAV315 (5’-TATATTTGGTGTGAAAGAAAGAAGCACATG −3’). Following transformation, clones were isolated and CRISPR arrays were screened using primers AV1171 and AV1015 to detect spacer deletions. Positive clones were then passaged without selection for two 1:100 dilutions and patched on plain and chloramphenicol plates to isolate clones that had lost the targeting plasmid.

### Phage construction

To create ΦAP1.1-spec, the K56 strain of *S. pyogenes* was transformed with pAV329, a plasmid containing spectinomycin resistance cassette and ~1 kilobase phage-homology arms, to insert the antibiotic resistance cassette immediately downstream the phage integrase in the lysogeny region. This strain was diluted 1:50 into 10 mls of BHI and supplemented with 5 mM CaCl_2_ and grown at 37 °C without agitation and infected with ΦAP1.1 at OD_600_ = 0.3. The resulting lysate was filtered through 0.45-μm syringe filters (Acrodisc) and used to infect K56. Five hours post infection cells were plated on 50 μg/ml spectinomycin. Insertion of the spectinomycin cassette into ΦAP1.1 was confirmed using primers AV1277 and AV1298.

Isolation of ΦAP1.1^*Δacr23*^ deletion was isolated by passaging ΦAP1.1-spec on a CEM strain containing a plasmid expressing a type III-A CRISPR-Cas system from *S. epidermidis* with a spacer targeting *acrIIA23* (5’-ATATTCCATTTTATTTCTCCTTATTACATAGATAA-3’, pAV449). 300 μl of overnight cultures of CEMΔΦ[Δ2-5] transformed with pAV392 mixed with 5 ml of 50% BHI agar supplemented with 5 mM CaCl_2_ and infected ΦAP1.1-spec. Escaper plaques were isolated and replaqued on this strain. Phage DNA was isolated and primers AV1339 and 1340 were used to amplify the targeted region. A phage was isolated with a deletion spanning nucleotide 37,881 to 39,044 of the ΦAP1.1-spec genome, encompassing *acrIIA23* and *orf1*.

Isolation of ΦAP1.1^Δ*acr23*-esc^ escaping from spacer 4 targeting of SF370 CRISPR array was carried out by passaging ΦAP1.1^Δ*acr23*^ on a CEM strain containing a plasmid expressing a type II-A CRISPR spacer (pAV392). Spacer 4 from SF370 contains a mismatch at position 8 when targeting the ΦAP1.1, which was corrected in the spacer cloned into pAV392 to perfectly target ΦAP1.1. 300 μl of overnight cultures of CEMΔΦ[Δ2-5] transformed with pAV392 mixed with 5 ml of 50% BHI agar supplemented with 5 mM CaCl_2_ and infected ΦAP1.1^Δ*acr23*^. Escaper plaques were isolated and replaqued on this strain. Phage DNA was isolated and primers AV1127 and AV1128 were used to amplify the spacer 4 target region and a phage with a G to A mutation in the first PAM position was isolated for subsequent experiments.

### Phage Sequencing

ΦAP1.1-spec DNA was purified as previously described (Varble et al., 2019). Phage DNA was sheared to 300 base pairs using the Covaris Ultrasonicator S220. Sheared DNA was prepped using the TruSeq Nano kit (Illumina) and and was sequenced using 150-nucleotide single-end read with an Illumina MiSeq. The phage genome was assembled using SPAdes (Bankevich et al., 2012) and the phage *cos* site was predicted using PhageTerm (Garneau et al., 2017).

### Quantification of Lysogens

To quantify lysogens, the indicated strain was diluted 1:30 into BHI supplemented with 5 mM CaCl_2_. At OD_600_ = 0.3, cultures were infected with 50 μl of ΦAP1.1-spec. Cells were harvested 90 minutes post infection and plated on BHI plates with 50 μg/ml spectinomycin. Individual colonies were restreaked and genomic DNA was isolated by resuspending colonies in colony lysis buffer (250 mM KCl, 5 mM MgCl_2_ 50 mM Tris-HCl at pH 9.0, 0.5% Triton X-100) supplemented with 500U PlyC (Nelson et al., 2006) and incubated at 37°C for 20 minutes, then 98°C for 10 minutes. Samples were centrifuged and supernatant was used for PCR amplification with primers H237 and AV1050 for the *attL* junction and AV1089 and AV1015 for the *attR* junction.

### Lysogen passaging

ΦAP1.1-spec lysogens integrated into the indicated direct repeat (DR) of CEM1ΔΦ were inoculated from single colonies and serially passaged by making 1:100 dilutions after 24 hours of growth. Half were passaged in the presence of a sub-lethal concentration of mitomycin C (0.1 μg/ml).

### Efficiency of plaquing assays

For *S. aureus*, phage titer assays were performed as previously described(Goldberg et al., 2014). Briefly, *S. aureus* strains with either the wildtype integrated type II-A locus from SF370 (JAV17) or the locus with a ΦNM4γ4 targeting spacer (JAV16, 5’-TGATACATTAACATTTAGTAAATCATTACG-3’) were transformed with either a Ctl vector (pJTR162), orf4 (pAV412), AcrIIA23 (pAV413), or orf1 (pAV414). Strains were outgrown for 2 hours. 30 minutes pre-collection aTc was added at a concentration of 12.5 ng/ml. 200 μl of cultures were mixed with 5 ml of 50% BHI agar supplemented with 5 mM CaCl_2_. Cells were plated on plain plates or plates supplemented with aTc. Serial dilutions of ΦNM4γ4 were then spotted on the plates.

### Cell Transformations

For *S. aureus*, transformations were carried out as previously described (Goldberg et al., 2014). For *S. pyogenes*, overnight cultures were diluted 1:50 into 250 mls of BHI and grown at 37 °C without agitation. At OD_600_ 0.3-0.4, cultures were pelleted and washed three times with cold 10% glycerol. Plasmid DNA was prepped from *Staphylococcus aureus* strain RN4220 (Kreiswirth et al., 1983), and dialyzed on 0.025 μm nitrocellulose filters (Millipore). 100 ng of DNA was added to 50 μl aliquots of electrocompetent cells and transformed in cold 1 mm cuvettes under the following settings: 2500 V, 25 μF, 600 Ω, 1 mm. After electroporation cells were resuspended in 950 μl of BHI and recovered at 37°C without agitation for two hours.

### Transformation Assays

For *S. pyogenes*, cells were prepped as described in cell transformation section. The plasmids used for targeting experiments were Ctl (pC194), Spacer 1 target (pAV344), Spacer 4 target as it appears in ΦAP1.1 (pAV354), Spacer 5 target (pAV346), and the three potential NCTC 13743 targets as they appear in ΦAP1.1 (pAV407). 100 ng of DNA was added to 50 μl aliquots of electrocompetent cells and transformed in cold 1 mm cuvettes under the following settings: 2500 V, 25 μF, 600 Ω, 1 mm. After electroporation cells were resuspended in 950 μl of BHI and recovered at 37°C without agitation for two hours. Serial dilutions were then made and plated on the appropriate antibiotics. For cultures where AcrIIA23 was expressed, cells transformed with Ctl (pLZ12spec) and AcrIIA23 (pAV432) were grown as above, with anhydrous tetracycline (aTc) added 30 minutes pre-collection at 12.5 ng/ml. Cells were treated as above and plated on the appropriate antibiotics plus or minus aTc.

### RNA isolation

Overnight cultures were diluted 1:50 into 100 mls of BHI and grown at 37 °C without agitation. At OD_600_ 0.3-0.4, cultures were pelleted. Cells were resuspended in 70 μl of PBS supplemented with 500U PlyC (Nelson et al., 2006) and incubated at 37°C for 30 minutes with shaking. 10 μl of 10% sarkosyl was then added and sample was then vortexed. RNA was then purified and DNase treated using isolated using the Direct-zol miniprep plus kit (Zymo Research) for RNA-seq or with TRIzol for Northern blots.

### Small RNA Northern blots

Northern blot analysis was carried out as previously described (Pall and Hamilton, 2008). 20 μg of purified RNA was loaded. AV1149 was used to blot for spacer 1, AV1152 for spacer 4, AV1153 for spacer 5, and AV1156 for 5S. Primers used in this study are detailed in Supplementary Table S2.

### RNA sequencing

For phage infection time courses, overnight cultures of *S. pyogenes* strain CEM were diluted 1:50 into 10 mls of BHI and grown at 37 °C without agitation. At OD_600_ 0.3 cells were infected with ΦAP1.1-spec and were pelleted and frozen at indicated time points. RNA was then isolated as described above. For lysogen sequencing an overnight culture of CEM with ΦAP1.1-spec integrated into direct repeat 3 was diluted 1:50 into 10 mls of BHI and grown at 37 °C without agitation. At OD_600_ 0.3, RNA was harvested as described as above. Ribosomal RNA was depleted using Ribo-Zero Plus rRNA Depletion Kit (Illumina) and subsequently purified using the RNA Clean and Concentrate −5 kit (Zymo Research). RNA was then prepped for deep-sequencing using the TruSeq Stranded mRNA Library Preparation Kit (Illumina) and run on the MiSeq System (Illumina). Reads were aligned to the ΦAP1.1-spec genome for the phage time course or the ΦAP1.1-spec genome integrated into direct repeat 3 of the CEM1ΔΦ CRISPR array using Bowtie 2 (Langmead and Salzberg, 2012). SAM files were normalized by total read count and converted to WIG files with a custom python script to plot coverage at each nucleotide. For the phage infection time course reads were mapped from nucleotide 35,754 to 39,363 of the ΦAP1.1-spec genome. For lysogen sequencing reads were mapped to this region plus the 286 base pairs of the 3’ portion of the CRISPR array.

### Characterizing phage *attP*

To isolate phage carrying spacers from the CEM1ΔΦ CRISPR array, overnight cultures of the indicated lysogens were diluted 1:50 into 10 mls of BHI and grown at 37 °C without agitation. At OD_600_ = 0.3 cells were induced with 0.5 μg/ml of mitomycin C. Four hours post-induction cells were spun down and supernatant was filtered through 0.45-μm syringe filters (Acrodisc). Phage DNA was purified from these inductions as previously described (Varble et al., 2019). From purified DNA, specific PCRs were carried out for spacer 1, 2 and 5. For spacers 1 and 2, AV1089 was used as the forward primer, while AV1149 and AV1151 were used as reverse primers for spacer 1 and 2 respectively. For spacer 5, H57 was the spacer 5-specfic forward primer, while AV1008 was the reverse primer. To deep sequence the phage *attP*, the phage junction was amplified from purified phage particles with AV1169 and AV1170, products above the typical 75-nucleotide phage *attP* were gel purified and prepped for deep-sequencing with the TruSeq Nano kit (Illumina). DNA was sequenced using 150-nucleotide single-end read with an Illumina MiSeq. A custom python script was used to quantify spacers in the phage *attP*, with read length limiting the detection of spacers in the *attP* to two.

### Detection of spacer transfer

Phage carrying spacers from the CEM1ΔΦ CRISPR array were isolated as described in the characterizing *attP* section. *S. pyogenes* strain K56 was diluted 1:30 into 10 mls of BHI and supplemented with 5 mM CaCl_2_ and grown at 37 °C without agitation. At OD_600_ = 0.3 cells were infected with phage induced from the indicated direct repeat. Cells were harvested five hours post infection. Genomic DNA was isolated using the Wizard^®^ Genomic DNA Purification kit (Promega). Cells were resuspended in 100 μl of 50mM EDTA with 500U PlyC and incubated at 37 °C with shaking for 30 minutes. 600 μl of Nuclei lysis solution was then added and DNA was purified following the manufacturer’s instructions. PCRs were then carried out to detect transfer of transfer of spacers either 5’ or 3’ of the integrated lysogen in K56. For the amplification of 5’ spacers, AV1171 was the forward primer, while AV1149 and AV1151 were used to specifically detect CEM1ΔΦ spacer’s one and three, respectively. For the amplification of 3’ spacers, AV1015 was the reverse primer, while H49 and H53 were used to specifically detect CEM1ΔΦ spacer’s one and three, respectively.

### Plasmid construction

Table S3 contains the information of the primers, templates and DNA fragments used to construct all the plasmids in this study.

### Next generation sequencing data

Next-generation sequencing data used to generate Figures 4A, 4B, 5D and 5E is deposited under BioProject accession number PRJNA668016.

## Supplementary Tables

**Supplementary table S1.** Strains used in this study.

| Strains | Description | Reference |
| --- | --- | --- |
| K56 | *S. pyogenes* strain | (Kjems, 1958) |
| SF370 | *S. pyogenes* strain with spacer targeting | (Ferretti et al., 2001) |
| CEM1ΔΦ | *S. pyogenes* strain, SF370 with no prophages | (Euler et al., 2016) |
| CEM1ΔΦ[Δ2-5] | *S. pyogenes* strain CEM1ΔΦ with spacers 2-5 deleted | This study |
| NCTC13743 | *S. pyogenes* strain | Accession no. SAMEA3879489 |
| NCTC13743[Δ1-10] | *S. pyogenes* strain NCTC 13743 with spacers 1-10 deleted | This study |
| RN4220 | *S. aureus* strain | (Kreiswirth et al., 1983) |
| JAV16 | *S. aureus* strain with integrated SF370 type II-A CRISPR targeting ΦNM4y4 | This study |
| JAV17 | *S. aureus* strain with integrated *S. pyogenes* type II-A CRISPR | This study |
| ΦAP1.1 | *S. pyogenes* phage | This study |
| ΦAP1.1-spec | ΦAP1.1 tagged with spectinomycin in lysogeny region | This study |
| ΦAP1.1<sup>Δacr23</sup> | ΦAP1.1-spec with *acrIIA23* deletion |  |
| ΦAP1.1<sup>Δacr23-esc</sup> | ΦAP1.1<sup>Δacr23</sup> escaper of spacer 4 from SF370 | This study |
| ΦNM4y4 | Obligately lytic *S. aureus* pac phage | (Heler et al., 2015) |
| pLZ12spec | Spectinomycin resistant cloning vector | (Husmann et al., 1995) |
| pC194 | Chloramphenicol-resistant *S. aureus* plasmid | (Horinouchi and Weisblum, 1982a) |
| pCasSA | *S. aureus* genome editing vector | (Chen et al., 2017) |
| pCL55 | Integration vector for *S. aureus* | (Luong and Lee, 2007) |
| pDB114 | Control spacer with *cas9* | (Bikard et al., 2014) |
| pE194 | Erythromycin-resistant *S. aureus* plasmid | (Horinouchi and Weisblum, 1982b) |
| pGG-BsaI-R | Type III-A CRISPR-Cas from *S. epidermidis* on pC194 | (Goldberg et al., 2014) |
| pJTR142 | Type III-A CRISPR-Cas from *S. epidermidis* on pLZ12spec | This study |
| pJTR162 | *S. aureus* tet inducible expression construct on pE194 | (Rostol and Marraffini, 2019) |
| pWJ40 | *S. pyogenes* type II-A CRISPR-Cas system on pC194 | (Heler et al., 2015) |
| pAV121 | Type II-A CRISPR-Cas from *S. pyogenes* on pCL55 | This study |
| pAV122 | Type II-A CRISPR-Cas from *S. pyogenes* on pCL55 targeting ΦNM4y4 | This study |
| pAV315 | Type II-A CRISPR-Cas from *S. pyogenes* targeting ΦAP1.1 on pC194 | This study |
| pAV329 | ΦAP1.1 editing plasmid to tag with spectinomycin | This study |
| pAV344 | Target site for SF370 spacer 1 on pC194 | This study |
| pAV346 | Target site from ΦAP1.1 for SF370 spacer 4 on pC194 | This study |
| pAV354 | Target site from for SF370 spacer 5 on pC194 | This study |
| pAV391 | Control sgRNA expressing vector on pC194 | This study |
| pAV392 | Perfect ΦAP1.1 spacer 4 sgRNA expressing vector on pC194 | This study |
| pAV407 | Target sites from ΦAP1.1 for NCTC 13743 spacers on pC194 | This study |
| pAV412 | ΦAP1.1 *orf4* overexpression plasmid on pE194 | This study |
| pAV413 | ΦAP1.1 *acrIIA23* overexpression plasmid on pE194 | This study |
| pAV414 | ΦAP1.1 *orf1* overexpression plasmid on pJTR162 | This study |
| pAV432 | ΦAP1.1 *acrIIA23* overexpression plasmid on pLZ12spec | This study |
| pAV449 | Type III-A *acrIIA23* targeting plasmid on pLZ12spec | This study |

**Supplementary table S2.**
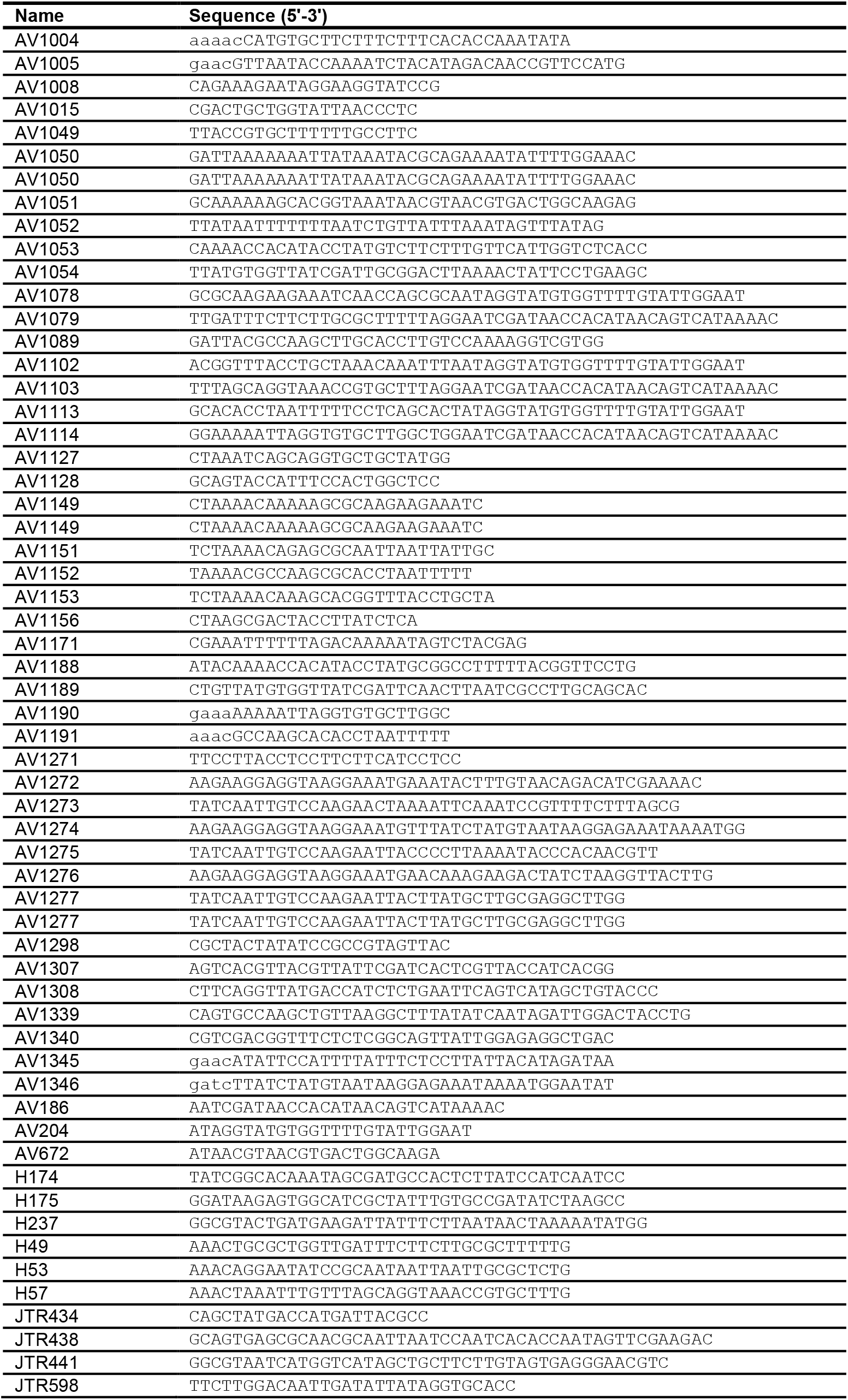
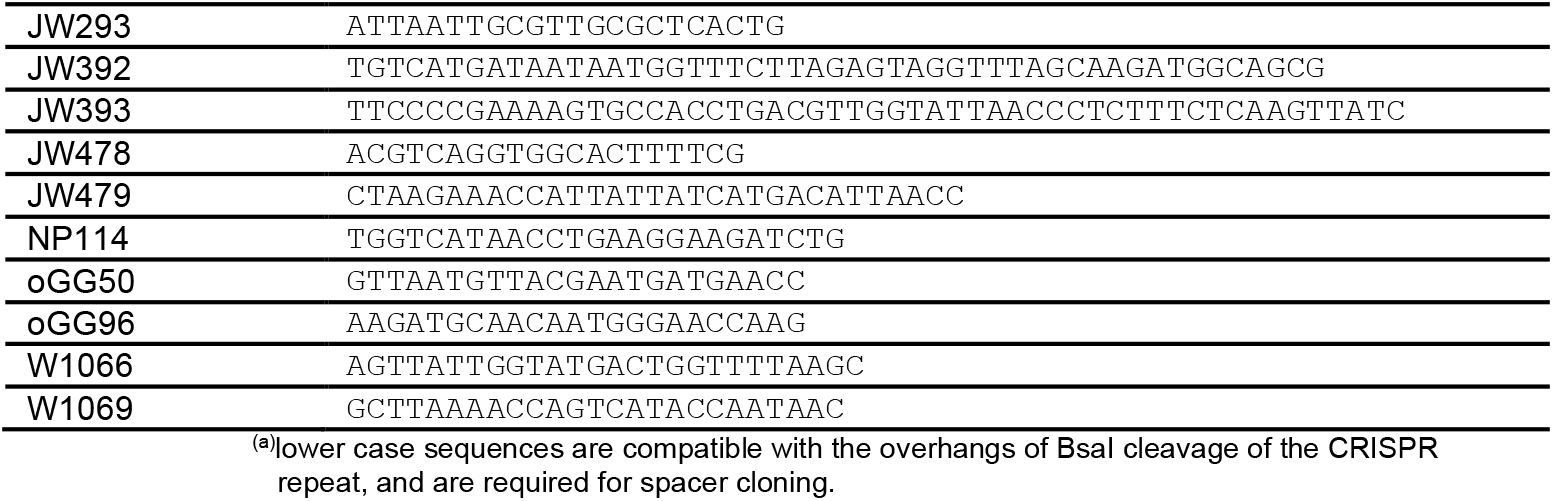
DNA oligonucleotides used in this study.

**Supplementary table S3.**
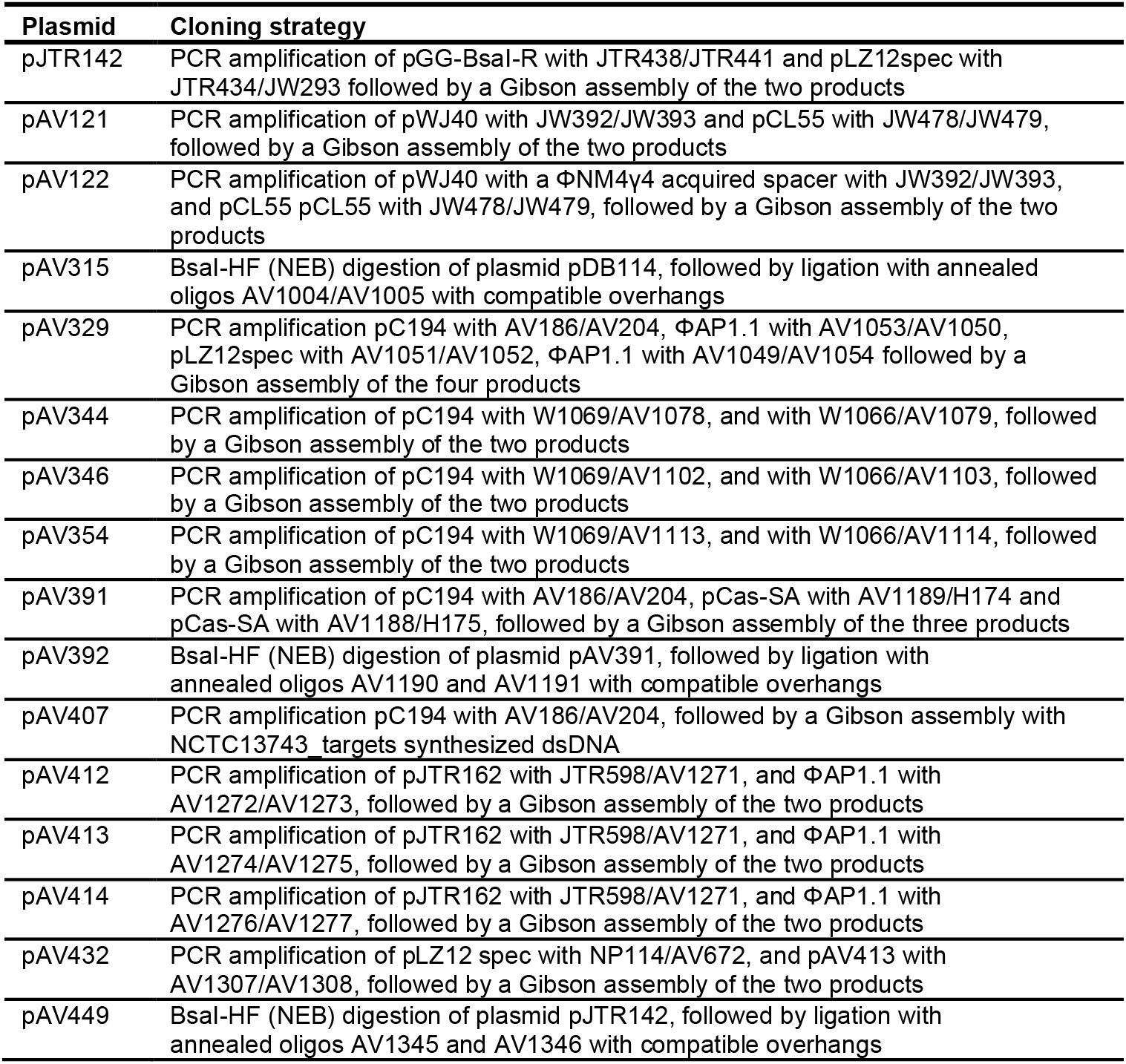
Plasmid construction in this study.

| Plasmid | Cloning strategy |
| --- | --- |
| pJTR142 | PCR amplification of pGG-BsaI-R with JTR438/JTR441 and pLZ12spec with JTR434/JW293 followed by a Gibson assembly of the two products |
| pAV121 | PCR amplification of pWJ40 with JW392/JW393 and pCL55 with JW478/JW479, followed by a Gibson assembly of the two products |
| pAV122 | PCR amplification of pWJ40 with a ΦNM4y4 acquired spacer with JW392/JW393, and pCL55 pCL55 with JW478/JW479, followed by a Gibson assembly of the two products |
| pAV315 | BsaI-HF (NEB) digestion of plasmid pDB114, followed by ligation with annealed oligos AV1004/AV1005 with compatible overhangs |
| pAV329 | PCR amplification pC194 with AV186/AV204, ΦAP1.1 with AV1053/AV1050, pLZ12spec with AV1051/AV1052, ΦAP1.1 with AV1049/AV1054 followed by a Gibson assembly of the four products |
| pAV344 | PCR amplification of pC194 with W1069/AV1078, and with W1066/AV1079, followed by a Gibson assembly of the two products |
| pAV346 | PCR amplification of pC194 with W1069/AV1102, and with W1066/AV1103, followed by a Gibson assembly of the two products |
| pAV354 | PCR amplification of pC194 with W1069/AV1113, and with W1066/AV1114, followed by a Gibson assembly of the two products |
| pAV391 | PCR amplification of pC194 with AV186/AV204, pCas-SA with AV1189/H174 and pCas-SA with AV1188/H175, followed by a Gibson assembly of the three products |
| pAV392 | BsaI-HF (NEB) digestion of plasmid pAV391, followed by ligation with annealed oligos AV1190 and AV1191 with compatible overhangs |
| pAV407 | PCR amplification pC194 with AV186/AV204, followed by a Gibson assembly with NCTC13743_targets synthesized dsDNA |
| pAV412 | PCR amplification of pJTR162 with JTR598/AV1271, and ΦAP1.1 with AV1272/AV1273, followed by a Gibson assembly of the two products |
| pAV413 | PCR amplification of pJTR162 with JTR598/AV1271, and ΦAP1.1 with AV1274/AV1275, followed by a Gibson assembly of the two products |
| pAV414 | PCR amplification of pJTR162 with JTR598/AV1271, and ΦAP1.1 with AV1276/AV1277, followed by a Gibson assembly of the two products |
| pAV432 | PCR amplification of pLZ12 spec with NP114/AV672, and pAV413 with AV1307/AV1308, followed by a Gibson assembly of the two products |
| pAV449 | BsaI-HF (NEB) digestion of plasmid pJTR142, followed by ligation with annealed oligos AV1345 and AV1346 with compatible overhangs |

### Synthetic DNA in this study

>NCTC13743_targets

~~~
ATACAAAACCACATACCTATAACATGTACGAGCTCAACAGTATTTTAGCAGATGTGTATGTCAAATTGATTGGTGGAGAGCCAGATGACCGGAACTA
TTGAGAGTCACAACATCAGGACTGGTCAATTAACTGTTGAGGAGTGGCAACGGCTTATCTATGATATGGGAGCAGGAACACCTTTAGGCAGTATGTT
TATCGAATTGGGTCTTGATACATCAAAATTTGATCCAATCGATAACCACATAACAG
~~~

## References

(2016). In Streptococcus pyogenes : Basic Biology to Clinical Manifestations, J.J. Ferretti, D.L. Stevens, and V.A. Fischetti, eds. (Oklahoma City (OK)).

Bankevich, A., Nurk, S., Antipov, D., Gurevich, A.A., Dvorkin, M., Kulikov, A.S., Lesin, V.M., Nikolenko, S.I., Pham, S., Prjibelski, A.D., et al. (2012). SPAdes: a new genome assembly algorithm and its applications to single-cell sequencing. J Comput Biol 19, 455–477.

Banks, D.J., Beres, S.B., and Musser, J.M. (2002). The fundamental contribution of phages to GAS evolution, genome diversification and strain emergence. Trends Microbiol 10, 515–521.

Barrangou, R., Fremaux, C., Deveau, H., Richards, M., Boyaval, P., Moineau, S., Romero, D.A., and Horvath, P. (2007). CRISPR provides acquired resistance against viruses in prokaryotes. Science 315, 1709–1712.

Bikard, D., Euler, C.W., Jiang, W., Nussenzweig, P.M., Goldberg, G.W., Duportet, X., Fischetti, V.A., and Marraffini, L.A. (2014). Exploiting CRISPR-Cas nucleases to produce sequence-specific antimicrobials. Nat Biotechnol 32, 1146–1150.

Bondy-Denomy, J., Davidson, A.R., Doudna, J.A., Fineran, P.C., Maxwell, K.L., Moineau, S., Peng, X., Sontheimer, E.J., and Wiedenheft, B. (2018). A Unified Resource for Tracking Anti-CRISPR Names. CRISPR J 1, 304–305.

Bondy-Denomy, J., Pawluk, A., Maxwell, K.L., and Davidson, A.R. (2013). Bacteriophage genes that inactivate the CRISPR/Cas bacterial immune system. Nature 493, 429–432.

Borges, A.L., Zhang, J.Y., Rollins, M.F., Osuna, B.A., Wiedenheft, B., and Bondy-Denomy, J. (2018). Bacteriophage Cooperation Suppresses CRISPR-Cas3 and Cas9 Immunity. Cell 174, 917–925 e910.

Brouns, S.J., Jore, M.M., Lundgren, M., Westra, E.R., Slijkhuis, R.J., Snijders, A.P., Dickman, M.J., Makarova, K.S., Koonin, E.V., and van der Oost, J. (2008). Small CRISPR RNAs guide antiviral defense in prokaryotes. Science 321, 960–964.

Chen, W., Zhang, Y., Yeo, W.S., Bae, T., and Ji, Q. (2017). Rapid and Efficient Genome Editing in Staphylococcus aureus by Using an Engineered CRISPR/Cas9 System. J Am Chem Soc 139, 3790–3795.

Coleman, D., Knights, J., Russell, R., Shanley, D., Birkbeck, T.H., Dougan, G., and Charles, I. (1991). Insertional inactivation of the Staphylococcus aureus beta-toxin by bacteriophage phi 13 occurs by site- and orientation-specific integration of the phi 13 genome. Mol Microbiol 5, 933–939.

Cong, L., Ran, F.A., Cox, D., Lin, S., Barretto, R., Habib, N., Hsu, P.D., Wu, X., Jiang, W., Marraffini, L.A., et al. (2013). Multiplex Genome Engineering Using CRISPR/Cas Systems. Science 339, 819–823.

Davidson, A.R., Lu, W.T., Stanley, S.Y., Wang, J., Mejdani, M., Trost, C.N., Hicks, B.T., Lee, J., and Sontheimer, E.J. (2020). Anti-CRISPRs: Protein Inhibitors of CRISPR-Cas Systems. Annu Rev Biochem 89, 309–332.

Deltcheva, E., Chylinski, K., Sharma, C.M., Gonzales, K., Chao, Y., Pirzada, Z.A., Eckert, M.R., Vogel, J., and Charpentier, E. (2011). CRISPR RNA maturation by trans-encoded small RNA and host factor RNase III. Nature 471, 602–607.

Desiere, F., McShan, W.M., van Sinderen, D., Ferretti, J.J., and Brussow, H. (2001). Comparative genomics reveals close genetic relationships between phages from dairy bacteria and pathogenic Streptococci: evolutionary implications for prophage-host interactions. Virology 288, 325–341.

Deveau, H., Barrangou, R., Garneau, J.E., Labonte, J., Fremaux, C., Boyaval, P., Romero, D.A., Horvath, P., and Moineau, S. (2008). Phage response to CRISPR-encoded resistance in *Streptococcus thermophilus*. J Bacteriol 190, 1390–1400.

Eitzinger, S., Asif, A., Watters, K.E., Iavarone, A.T., Knott, G.J., Doudna, J.A., and Minhas, F. (2020). Machine learning predicts new anti-CRISPR proteins. Nucleic Acids Res 48, 4698–4708.

Euler, C.W., Juncosa, B., Ryan, P.A., Deutsch, D.R., McShan, W.M., and Fischetti, V.A. (2016). Targeted Curing of All Lysogenic Bacteriophage from Streptococcus pyogenes Using a Novel Counter-selection Technique. PLoS One 11, e0146408.

Ferretti, J.J., McShan, W.M., Ajdic, D., Savic, D.J., Savic, G., Lyon, K., Primeaux, C., Sezate, S., Suvorov, A.N., Kenton, S., et al. (2001). Complete genome sequence of an M1 strain of *Streptococcus pyogenes*. Proc Natl Acad Sci USA 98, 4658–4663.

Fiebig, A., Loof, T.G., Babbar, A., Itzek, A., Koehorst, J.J., Schaap, P.J., and Nitsche-Schmitz, D.P. (2015). Comparative genomics of Streptococcus pyogenes M1 isolates differing in virulence and propensity to cause systemic infection in mice. Int J Med Microbiol 305, 532–543.

Fischetti, V.A. (2007). In vivo acquisition of prophage in Streptococcus pyogenes. Trends Microbiol 15, 297–300.

Forsberg, K.J., Bhatt, I.V., Schmidtke, D.T., Javanmardi, K., Dillard, K.E., Stoddard, B.L., Finkelstein, I.J., Kaiser, B.K., and Malik, H.S. (2019). Functional metagenomics-guided discovery of potent Cas9 inhibitors in the human microbiome. Elife 8.

Garneau, J.E., Dupuis, M.E., Villion, M., Romero, D.A., Barrangou, R., Boyaval, P., Fremaux, C., Horvath, P., Magadan, A.H., and Moineau, S. (2010). The CRISPR/Cas bacterial immune system cleaves bacteriophage and plasmid DNA. Nature 468, 67–71.

Garneau, J.R., Depardieu, F., Fortier, L.C., Bikard, D., and Monot, M. (2017). PhageTerm: a tool for fast and accurate determination of phage termini and packaging mechanism using next-generation sequencing data. Sci Rep 7, 8292.

Gasiunas, G., Barrangou, R., Horvath, P., and Siksnys, V. (2012). Cas9-crRNA ribonucleoprotein complex mediates specific DNA cleavage for adaptive immunity in bacteria. Proc Natl Acad Sci USA 109, E2579–2586.

Goldberg, G.W., Jiang, W., Bikard, D., and Marraffini, L.A. (2014). Conditional tolerance of temperate phages via transcription-dependent CRISPR-Cas targeting. Nature 514, 633–637.

Heler, R., Samai, P., Modell, J.W., Weiner, C., Goldberg, G.W., Bikard, D., and Marraffini, L.A. (2015). Cas9 specifies functional viral targets during CRISPR-Cas adaptation. Nature 519, 199–202.

Horinouchi, S., and Weisblum, B. (1982a). Nucleotide sequence and functional map of pC194, a plasmid that specifies inducible chloramphenicol resistance. J Bacteriol 150, 815–825.

Horinouchi, S., and Weisblum, B. (1982b). Nucleotide sequence and functional map of pE194, a plasmid that specifies inducible resistance to macrolide, lincosamide, and streptogramin type B antibodies. J Bacteriol 150, 804–814.

Howard-Varona, C., Hargreaves, K.R., Abedon, S.T., and Sullivan, M.B. (2017). Lysogeny in nature: mechanisms, impact and ecology of temperate phages. ISME J 11, 1511–1520.

Husmann, L.K., Scott, J.R., Lindahl, G., and Stenberg, L. (1995). Expression of the Arp protein, a member of the M protein family, is not sufficient to inhibit phagocytosis of *Streptococcus pyogenes*. Infect Immun 63, 345–348.

Hynes, A.P., Rousseau, G.M., Lemay, M.L., Horvath, P., Romero, D.A., Fremaux, C., and Moineau, S. (2017). An anti-CRISPR from a virulent streptococcal phage inhibits Streptococcus pyogenes Cas9. Nat Microbiol 2, 1374–1380.

Jansen, R., Embden, J.D., Gaastra, W., and Schouls, L.M. (2002). Identification of genes that are associated with DNA repeats in prokaryotes. Mol Microbiol 43, 1565–1575.

Jinek, M., Chylinski, K., Fonfara, I., Hauer, M., Doudna, J.A., and Charpentier, E. (2012). A programmable dual-RNA-guided DNA endonuclease in adaptive bacterial immunity. Science 337, 816–821.

Kjems, E. (1958). Studies on streptococcal bacteriophages. 2. Adsorption, lysogenization, and one-step growth experiments. Acta Pathol Microbiol Scand 42, 56–66.

Kreiswirth, B.N., Lofdahl, S., Betley, M.J., O’Reilly, M., Schlievert, P.M., Bergdoll, M.S., and Novick, R.P. (1983). The toxic shock syndrome exotoxin structural gene is not detectably transmitted by a prophage. Nature 305, 709–712.

Landsberger, M., Gandon, S., Meaden, S., Rollie, C., Chevallereau, A., Chabas, H., Buckling, A., Westra, E.R., and van Houte, S. (2018). Anti-CRISPR Phages Cooperate to Overcome CRISPR-Cas Immunity. Cell 174, 908–916 e912.

Langmead, B., and Salzberg, S.L. (2012). Fast gapped-read alignment with Bowtie 2. Nat Methods 9, 357–359.

Lee, C.Y., Buranen, S.L., and Ye, Z.H. (1991). Construction of single-copy integration vectors for *Staphylococcus aureus*. Gene 103, 101–105.

Lee, C.Y., and Iandolo, J.J. (1986). Lysogenic conversion of staphylococcal lipase is caused by insertion of the bacteriophage L54a genome into the lipase structural gene. J Bacteriol 166, 385–391.

Luong, T.T., and Lee, C.Y. (2007). Improved single-copy integration vectors for *Staphylococcus aureus*. J Microbiol Methods 70, 186–190.

Mahendra, C., Christie, K.A., Osuna, B.A., Pinilla-Redondo, R., Kleinstiver, B.P., and Bondy-Denomy, J. (2020). Broad-spectrum anti-CRISPR proteins facilitate horizontal gene transfer. Nat Microbiol 5, 620–629.

Makarova, K.S., Wolf, Y.I., Iranzo, J., Shmakov, S.A., Alkhnbashi, O.S., Brouns, S.J.J., Charpentier, E., Cheng, D., Haft, D.H., Horvath, P., et al. (2020). Evolutionary classification of CRISPR-Cas systems: a burst of class 2 and derived variants. Nat Rev Microbiol 18, 67–83.

Mali, P., Yang, L., Esvelt, K.M., Aach, J., Guell, M., Dicarlo, J.E., Norville, J.E., and Church, G.M. (2013). RNA-Guided Human Genome Engineering via Cas9. Science 339, 823–826.

Marino, N.D., Zhang, J.Y., Borges, A.L., Sousa, A.A., Leon, L.M., Rauch, B.J., Walton, R.T., Berry, J.D., Joung, J.K., Kleinstiver, B.P., et al. (2018). Discovery of widespread type I and type V CRISPR-Cas inhibitors. Science 362, 240–242.

Marraffini, L.A., and Sontheimer, E.J. (2008). CRISPR interference limits horizontal gene transfer in staphylococci by targeting DNA. Science 322, 1843–1845.

McCullor, K., Postoak, B., Rahman, M., King, C., and McShan, W.M. (2018). Genomic Sequencing of High-Efficiency Transducing Streptococcal Bacteriophage A25: Consequences of Escape from Lysogeny. J Bacteriol 200.

McGinn, J., and Marraffini, L.A. (2016). CRISPR-Cas systems optimize their immune response by specifying the site of spacer integration. Mol Cell 64, 616–623.

Modell, J.W., Jiang, W., and Marraffini, L.A. (2017). CRISPR-Cas systems exploit viral DNA injection to establish and maintain adaptive immunity. Nature 544, 101–104.

Nelson, D., Schuch, R., Chahales, P., Zhu, S., and Fischetti, V.A. (2006). PlyC: a multimeric bacteriophage lysin. Proc Natl Acad Sci U S A 103, 10765–10770.

Nozawa, T., Furukawa, N., Aikawa, C., Watanabe, T., Haobam, B., Kurokawa, K., Maruyama, F., and Nakagawa, I. (2011). CRISPR inhibition of prophage acquisition in *Streptococcus pyogenes*. PLoS One 6, e19543.

Nussenzweig, P.M., McGinn, J., and Marraffini, L.A. (2019). Cas9 Cleavage of Viral Genomes Primes the Acquisition of New Immunological Memories. Cell Host Microbe 26, 515–526 e516.

Osuna, B.A., Karambelkar, S., Mahendra, C., Sarbach, A., Johnson, M.C., Kilcher, S., and Bondy-Denomy, J. (2020). Critical Anti-CRISPR Locus Repression by a Bi-functional Cas9 Inhibitor. Cell Host Microbe.

Pall, G.S., and Hamilton, A.J. (2008). Improved northern blot method for enhanced detection of small RNA. Nat Protoc 3, 1077–1084.

Pawluk, A., Staals, R.H., Taylor, C., Watson, B.N., Saha, S., Fineran, P.C., Maxwell, K.L., and Davidson, A.R. (2016). Inactivation of CRISPR-Cas systems by anti-CRISPR proteins in diverse bacterial species. Nat Microbiol 1, 16085.

Perez-Casal, J., Caparon, M.G., and Scott, J.R. (1991). Mry, a trans-acting positive regulator of the M protein gene of *Streptococcus pyogenes* with similarity to the receptor proteins of two-component regulatory systems. J Bacteriol 173, 2617–2624.

Rauch, B.J., Silvis, M.R., Hultquist, J.F., Waters, C.S., McGregor, M.J., Krogan, N.J., and Bondy-Denomy, J. (2017). Inhibition of CRISPR-Cas9 with Bacteriophage Proteins. Cell 168, 150–158 e110.

Rostol, J.T., and Marraffini, L.A. (2019). Non-specific degradation of transcripts promotes plasmid clearance during type III-A CRISPR-Cas immunity. Nat Microbiol 4, 656–662.

Stanley, S.Y., Borges, A.L., Chen, K.H., Swaney, D.L., Krogan, N.J., Bondy-Denomy, J., and Davidson, A.R. (2019). Anti-CRISPR-Associated Proteins Are Crucial Repressors of Anti-CRISPR Transcription. Cell 178, 1452–1464 e1413.

Taylor, A.L. (1963). Bacteriophage-Induced Mutation in Escherichia Coli. Proc Natl Acad Sci U S A 50, 1043–1051.

Uribe, R.V., van der Helm, E., Misiakou, M.A., Lee, S.W., Kol, S., and Sommer, M.O.A. (2019). Discovery and Characterization of Cas9 Inhibitors Disseminated across Seven Bacterial Phyla. Cell Host Microbe 25, 233–241 e235.

Varble, A., Meaden, S., Barrangou, R., Westra, E.R., and Marraffini, L.A. (2019). Recombination between phages and CRISPR-cas loci facilitates horizontal gene transfer in staphylococci. Nat Microbiol 4, 956–963.

Watson, B.N.J., Staals, R.H.J., and Fineran, P.C. (2018). CRISPR-Cas-Mediated Phage Resistance Enhances Horizontal Gene Transfer by Transduction. MBio 9.

Wright, A.V., and Doudna, J.A. (2016). Protecting genome integrity during CRISPR immune adaptation. Nat Struct Mol Biol 23, 876–883.

Xiao, Y., Ng, S., Nam, K.H., and Ke, A. (2017). How type II CRISPR-Cas establish immunity through Cas1-Cas2-mediated spacer integration. Nature 550, 137–141.

Yamada, S., Shibasaki, M., Murase, K., Watanabe, T., Aikawa, C., Nozawa, T., and Nakagawa, I. (2019). Phylogenetic relationship of prophages is affected by CRISPR selection in Group A Streptococcus. BMC Microbiol 19, 24.

Yin, Y., Yang, B., and Entwistle, S. (2019). Bioinformatics Identification of Anti-CRISPR Loci by Using Homology, Guilt-by-Association, and CRISPR Self-Targeting Spacer Approaches. mSystems 4.

